# Effect of pregnancy and hypertension on kidney function in female rats: Modeling and functional implications

**DOI:** 10.1101/2022.12.15.520674

**Authors:** Melissa M. Stadt, Crystal A. West, Anita T. Layton

## Abstract

Throughout pregnancy, the kidneys undergo significant adaptations in morphology, hemodynamics, and transport to achieve the volume and electrolyte retention required to support a healthy pregnancy. Additionally, during pregnancies complicated by chronic hypertension, altered renal function from normal pregnancy occurs. The goal of this study is to analyze how inhibition of critical transporters affects gestational kidney function as well as how renal function is affected during chronic hypertension in pregnancy. To do this, we developed epithelial cell-based multi-nephron computational models of solute and water transport in the kidneys of a female rat in mid- and late pregnancy. We simulated the effects of key individual pregnancy-induced changes on renal Na^+^ and K^+^ transport: proximal tubule length, Na^+^/H^+^ exchanger isoform 3 (NHE3) activity, epithelial Na^+^ channel activity (ENaC), K^+^ secretory channel expression, and H^+^-K^+^-ATPase activity. Additionally, we conducted simulations to predict the effects of inhibition and knockout of the ENaC and H^+^-K^+^-ATPase transporters on virgin and pregnant rat kidneys. Our simulation results predicted that the ENaC and H^+^-K^+^-ATPase transporters are essential for sufficient Na^+^ and K^+^ reabsorption during pregnancy. Last, we developed models to capture changes made during hypertension in female rats and considered what may occur when a rat with chronic hypertension becomes pregnant. Model simulations predicted that in hypertension for a pregnant rat there is a similar shift in Na^+^ transport from the proximal tubules to the distal tubules as in a virgin rat.

## Introduction

Normal pregnancy is characterized by complicated and multifactorial adaptations in the maternal body in almost all physiological systems (1,2). These changes are dynamic, often competing, and can be counterintuitive when considered in isolation. One example is the significant rise in plasma volume, which if considered independently, would increase blood pressure to dangerous levels. However, in a normal pregnancy, blood pressure typically decreases due to changes in the cardiovascular system (3).

During a healthy pregnancy, a new organ, the placenta, develops and requires adequate blood flow to support sufficient oxygenation, nutrition, and maturation of the fetus (4). Notably, the placenta is also an endocrine organ which secretes hormones that alter maternal body function (1,4). By the end of gestation, progesterone levels rise approximately 100-fold and estrogen levels by about 50-fold (5), which can significantly impact many physiological systems. Their substantial increase can largely be attributed to placental secretion (1,2,4). Remarkably, the maternal body adapts to support the nutrient and electrolyte requirements of the rapidly growing fetus and placenta, as well as normal tissue and organ perfusion. Indeed, when the maternal body does not sufficiently adapt, dangerous gestational disorders, or fetal growth restriction may occur (6–9).

Chronic hypertension is one of the most common medical disorders globally, affecting 45% of all U.S. adults and an estimated 21% of women of reproductive age (10–12). In pregnancy, chronic hypertension is defined as hypertension that is diagnosed before or during the first 20 weeks of gestation (10,13,14). Interestingly, due to normal pregnancy-induced vascular adaptations, oftentimes women with chronic hypertension experience a decrease in blood pressure during pregnancy, sometimes being able to decrease medications or even become normotensive (13). Pregnancy has also been shown to be antihypertensive for spontaneously hypertensive rats (15,16).

While many women who have chronic hypertension are able to sustain a healthy pregnancy, they do have increased risk of dangerous adverse health effects such as superimposed preeclampsia, fetal growth restriction, preterm birth, and placental abruption (10,13,17). For example, preeclampsia, which is one of the most dangerous complications of pregnancy, occurs in about 17-25% of pregnancies in women with chronic hypertension compared with 3-5% of pregnancies in the general population (13,18). Additionally, pregnancies complicated by chronic hypertension have been linked to adverse health effects on the child throughout their lifetime (10,19,20). The prevalence chronic hypertension in pregnant women has doubled over the last decade, and this trend is expected to continue as average maternal age and obesity rates increase among women of childbearing age (10,14,21). Indeed, it is important to have a full understanding of the impacts of chronic hypertension and pregnancy, not only for the perinatal health of the fetus and mother, but also to prevent impacts of future disease caused by prenatal insults.

The kidneys play an essential role in regulating body homeostasis, namely water, electrolyte, and acid-base balance (22–25). During pregnancy, the plasma volume of the mother must expand drastically to support the rapidly developing fetus and placenta (3). In a virgin or non-pregnant rat, almost all Na^+^ intake is excreted through urine, but in pregnancy this is not the case: there is net Na^+^ retention in a pregnant rat (26,27). Additionally, in late pregnancy, maternal net K^+^ retention occurs (26–29). These changes are supported by the kidneys via major adaptations that are made in renal hemodynamics (30–33), morphology (30,34), and nephron transport (26,27). Specifically, this includes increased filtration to the kidneys, increased kidney volume, and altered transporter activity along the various nephron segments.

The kidneys play an essential role in long-term blood pressure control. Altered renal function in a rat occurs during hypertension (35–38). Specifically, in female rats, chronic hypertension has been shown to decrease Na^+^ transporter activity in the proximal segments as well as increase Na^+^ transporter activity in the distal segments (37). In this paper, we use computational modeling to predict how chronic hypertension impacts renal transport along the nephrons in a virgin female rat and pregnant rat.

In this study, we developed computational models of a full kidney in a pregnant rat to investigate the impact of key *pregnancy-induced* renal adaptations on electrolyte handling *in silico.* We built two computational models: one that represents kidney function at mid-pregnancy and another that represents kidney function at late pregnancy. The female-specific rat kidney model developed by Hu et al. (39) is used as a virgin control in this study. Two separate models for mid- and late pregnancy stages were necessary because of the changing demands of a growing fetus and placenta through gestation. Using these models, we investigated the impact of key pregnancy-induced renal adaptations on Na^+^ and K^+^ handling in a full kidney. Additionally, we considered the impact of epithelial Na^+^ channel and H^+^-K^+^-ATPase knockout and inhibition on renal Na^+^ and K^+^ transport during pregnancy based on existing experimental studies (Refs. (40,41)). Finally, we extend the models to investigate how hypertension may affect kidney function in a virgin rat and during pregnancy. The objective of this study is to answer the following questions: (1) To what extent can individual *pregnancy-induced* renal adaptations impact renal Na^+^ and K^+^ handling? (2) How does distal segment transporter inhibition and knockout impact fine tuning of Na^+^ and K^+^ during pregnancy? (3) What differences in renal electrolyte handling are predicted when a hypertensive female rat becomes pregnant?

## Materials and Methods

In our previous study (Ref. (27)), we developed epithelial cell-based models of a superficial nephron in the kidney of a virgin, mid-pregnant (MP), and late pregnant (LP) rat. However, the superficial nephrons only make up about 2/3 of the total nephron population, while the remaining nephrons are juxtamedullary nephrons, whose loops of Henle reach into the inner medulla. Hence in this study, we aimed to capture full kidney function by extending the models to represent both superficial and juxtamedullary nephrons. The resulting multi-nephron models typically yield more realistic predictions of overall kidney function and have been extensively developed and used to study kidney function in non-pregnant rats (39,42,43). To investigate the impact of pregnancy on kidney function, we developed *pregnancy-specific multi-nephron models* in MP and LP rats. Additionally, we developed a multi-nephron model of a hypertensive virgin (female) rat and considered what may happen in the kidneys when a hypertensive female becomes pregnant. Note that the original nephron transport model equations (see Refs. (39,42,43)) are based on mass conservation which are also valid in pregnancy and hypertension. Hence, those same equations are used in the MP, LP, and hypertensive models, but appropriate parameter values are changed to account for the appropriate renal adaptations.

### Multi-nephron epithelial transport model

Each nephron segment is modeled as a tubule lined by a layer of epithelial cells in which the apical and basolateral transporters vary depending on the cell type (i.e., segment, which part of segment, intercalated and principal cells; see Figure 1). The model accounts for the following 15 solutes: Na^+^, K^+^, Cl^-^, HCO_3_^-^, H_2_CO_3_, CO_2_, NH_3_, NH_4_^+^, HPO_4_^2-^, H_2_PO_4_^-^, H^+^, HCO_2_^-^, H_2_CO_2_, urea, and glucose. The model consists of a large system of coupled ordinary differential and algebraic equations, solved for steady state, and predicts luminal fluid flow, hydrostatic pressure, membrane potential, luminal and cytosolic solute concentrations, and transcellular and paracellular fluxes through transporters and channels. A schematic diagram of the various cell types, with MP, LP, and hypertension changes highlighted, is given in Figure 1.

**Figure 1.**
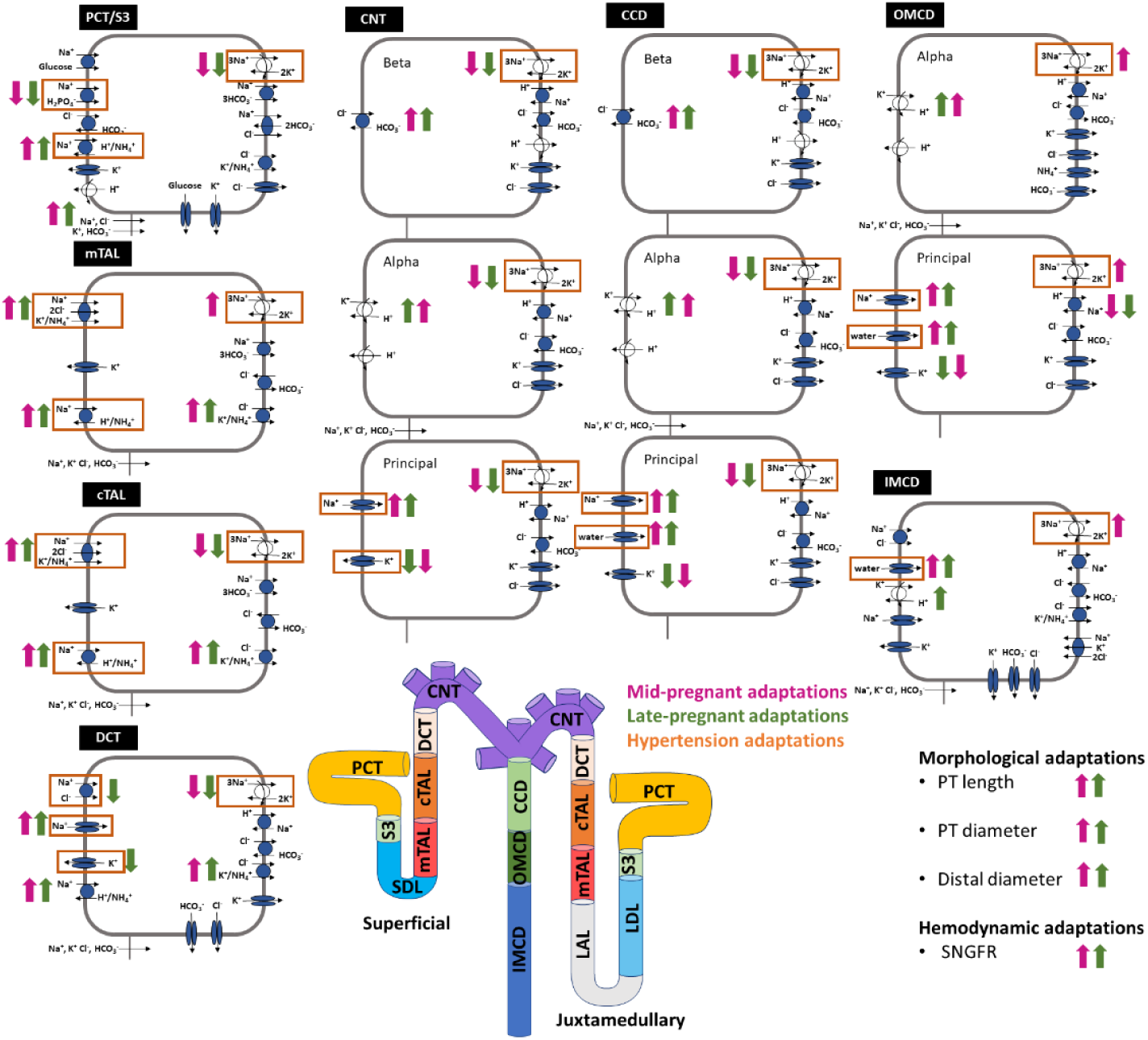
Schematic diagram of the nephrons represented in the models. The model includes a superficial nephron and five representative juxtamedullary nephrons scaled by the appropriate population ratio. Here one superficial and one juxtamedullary nephron are illustrated which come together to meet in the collecting duct. The cell diagrams represent cells from the various nephron segments and display the main Na^+^, K^+^, and Cl^-^ transporters as well as aquaporin water channels. Juxtamedullary descending thin limb (LDL) and thin ascending limb (LAL) cells are not shown and are represented by single-barrier transport models (see Ref. (43) for details). Mid-pregnant adaptations from virgin values are indicated by pink arrows. Late-pregnant adaptations from virgin values are indicated by green arrows. Upward orientation indicates that the transporter is increased in the respective model. Details are shown in Table 1. Transporters that are changed in the hypertensive models are boxed in orange. Details are shown in Table 3. CCD, cortical collecting duct; cTAL, cortical thick ascending limb; CNT, connecting tubule; DCT, distal convoluted tubule; IMCD, inner medullary collecting duct; LAL, thin ascending limb; LDL, thin descending limb; mTAL, medullary thick ascending limb; OMCD, outer medullary collecting duct; PCT, proximal convoluted tubule; SDL, short descending limb; SNGFR, single-nephron glomerular filtration rate; S3, proximal straight tubule. Cell and nephron diagram adapted from Refs. (27,43).

### Superficial vs. juxtamedullary nephrons

Kidneys have multiple types of nephrons; superficial and juxtamedullary nephrons make up most of them (43–46). As noted above, in a rat, about two-thirds of the nephron population are superficial nephrons: the remaining third are called juxtamedullary nephrons. The two nephron types differ in several key ways. The first major difference is that the juxtamedullary nephrons have loops of Henle that reach into the inner medulla. Specifically, the superficial nephron models include the proximal, short descending limb, thick ascending limb, distal convoluted tubule, connecting tubule, and the collecting duct. The juxtamedullary nephron models include all the same segments of the superficial nephron with the addition of the long descending limbs and ascending thin limbs; these are the segments of the loops of Henle that extend into the inner medulla (see Figure 1). The second major difference is that filtration to the juxtamedullary nephrons is higher than the filtered load delivered to the superficial nephrons. More details about the superficial versus juxtamedullary nephrons and how they are modeled is given in the *Supplementary Materials* and in Ref. (43).

### Pregnancy-specific models

We created pregnancy-specific models to simulate kidney function in MP and LP by using the virgin (female-specific) multiple nephron epithelial transport model developed in Ref. (39) and increasing or decreasing relevant virgin model parameter values based on experimental findings in the literature. The specific MP-to-virgin and LP-to-virgin parameter value ratios are listed in Table 1.

**Table 1.** MP-to-virgin and LP-to-virgin ratios of parameter values used in the MP and LP models, respectively. Virgin model parameter values were used from Ref. (39) and changed by the respective ratio for the MP and LP models. Parameter values for the MP and LP models were chosen based on experimental literature and are discussed in Ref. (27). Distal segments refers to all segments after the PCT and S3 segments (i.e., after the PT). Ratios are from the respective MP or LP model to the virgin model parameter value. The ratios are the same for the juxtamedullary and superficial nephrons. PCT, proximal convoluted tubule; S3, proximal straight tubule; SDL, short descending limb; LDL, thin descending limb; LAL, thin ascending limb; mTAL, medullary thick ascending limb; cTAL, cortical thick ascending limb; DCT, distal convoluted tubule; CNT, connecting tubule; CCD, cortical collecting duct; OMCD, outer medullary collecting duct; IMCD, inner medullary collecting duct; PT, proximal tubule; P_f_, water permeability; P_K_, K^+^ permeability; NHE3, Na^+^/H^+^ exchanger isoform 3; ENaC, epithelial Na^+^ channel; NKCC2, Na^+^-K^+^-2Cl^-^ cotransporter 2; NaPi2, Na^+^-H_2_PO_4_^-^ cotransporter 2; NCC, Na^+^-Cl^-^ cotransporter; KCC, K^+^-Cl^-^ cotransporter.

| Parameter | Ratios |  |
| --- | --- | --- |
|  | MP-to-virgin | LP-to-virgin |
| **Hemodynamics** |  |  |
| SNGFR | 1.30 | 1.20 |
| **Plasma Concentration** |  |  |
| $Na^+$ | 0.95 | 0.95 |
| $K^+$ | 1.00 | 1.15 |
| $Cl^-$ | 0.95 | 0.95 |
| $HCO_3^-$ | 0.95 | 0.95 |
| **PCT** |  |  |
| NHE3 activity | 1.40 | 1.20 |
| $Na^+-K^+$-ATPase activity | 0.75 | 0.65 |
| $P_f$ (transcellular) | 1.00 | 1.00 |
| KCC cotransporter | 1.35 | 1.30 |
| NaPi2 | 0.90 | 0.85 |
| **S3** |  |  |
| NHE3 activity | 1.40 | 1.20 |
| $Na^+-K^+$-ATPase activity | 0.73 | 0.65 |
| $P_f$ (transcellular) | 1.00 | 1.00 |
| KCC cotransporter | 1.35 | 1.35 |
| NaPi2 | 0.90 | 0.85 |
| **SDL** |  |  |
| $P_f$ (transcellular) | 1.10 | 1.50 |
| **LDL** |  |  |
| $P_f$ (transcellular) | 1.10 | 1.50 |
| **mTAL** |  |  |
| NKCC2 activity | 1.16 | 1.50 |
| NHE3 activity | 1.20 | 1.18 |
| KCC4 activity | 1.40 | 1.30 |
| Na<sup>+</sup>-K<sup>+</sup>-ATPase activity | 1.20 | 1.00 |
| **cTAL** |  |  |
| NKCC2 activity | 1.16 | 1.50 |
| NHE3 activity | 1.20 | 1.18 |
| KCC4 activity | 1.35 | 1.30 |
| Na<sup>+</sup>-K<sup>+</sup>-ATPase activity | 0.73 | 0.65 |
| **DCT** |  |  |
| P<sub>K</sub> (apical, DCT2 only) | 0.65 | 0.55 |
| Na<sup>+</sup>-K<sup>+</sup>-ATPase activity | 0.80 | 0.75 |
| NHE2 activity | 1.20 | 1.18 |
| KCC activity | 1.40 | 1.30 |
| NCC activity | 1.00 | 0.45 |
| ENaC activity | 2.08 | 2.15 |
| **CNT** |  |  |
| P<sub>K</sub> (apical) | 0.65 | 0.55 |
| Na<sup>+</sup>-K<sup>+</sup>-ATPase activity | 0.80 | 0.78 |
| H<sup>+</sup>-K<sup>+</sup>-ATPase activity | 2.25 | 2.75 |
| H<sup>+</sup>-ATPase activity | 1.00 | 1.00 |
| Pendrin activity | 1.65 | 1.70 |
| ENaC activity | 2.08 | 2.15 |
| **CCD** |  |  |
| P<sub>f</sub> (transcellular) | 1.40 | 1.40 |
| P<sub>K</sub> (apical) | 0.78 | 0.75 |
| Na<sup>+</sup>-K<sup>+</sup>-ATPase activity | 0.80 | 0.75 |
| H<sup>+</sup>-K<sup>+</sup>-ATPase activity | 2.25 | 2.75 |
| H<sup>+</sup>-ATPase activity | 1.00 | 1.00 |
| Pendrin activity | 1.65 | 1.70 |
| ENaC activity | 2.08 | 2.15 |
| **OMCD** |  |  |
| P<sub>f</sub> (transcellular) | 1.40 | 1.40 |
| P<sub>K</sub> (apical) | 0.88 | 0.85 |
| Na<sup>+</sup>-K<sup>+</sup>-ATPase activity | 1.20 | 1.08 |
| H<sup>+</sup>-ATPase activity | 1.00 | 1.00 |
| H<sup>+</sup>-K<sup>+</sup>-ATPase activity | 2.25 | 2.75 |
| ENaC activity | 2.08 | 2.15 |
| Na<sup>+</sup>-Cl<sup>-</sup> cotransporter | 1.15 | 1.00 |
| NHE1 activity | 0.90 | 0.90 |
| **IMCD** |  |  |
| P<sub>f</sub> (transcellular) | 1.80 | 1.80 |
| NKCC1 activity | 1.00 | 1.00 |
| Na<sup>+</sup>-K<sup>+</sup>-ATPase activity | 1.20 | 1.08 |
| KCC cotransporter | 1.40 | 1.30 |
| Na<sup>+</sup>-Cl<sup>-</sup> cotransporter | 1.15 | 1.00 |
| H<sup>+</sup>-K<sup>+</sup>-ATPase activity | 2.25 | 2.75 |
| **Morphology** |  |  |
| PT length | 1.15 | 1.17 |
| PT diameter | 1.07 | 1.07 |
| Distal length | 1.00 | 1.00 |
| Distal diameter | 1.05 | 1.05 |

In the virgin model, the superficial SNGFR is 24 nL/min and the juxtamedullary SNGFR is 33.6 nL/min (39). During pregnancy, there is a significant increase in GFR compared to the virgin values (3,32,33). Specifically, in rats, the GFR is increased by about 30% in MP and drops in LP to 20% above pre-gestational (virgin) GFR (3,32,33). The GFR is the total filtration by a kidney, hence is a total combination of the SNGFR of the superficial and juxtamedullary nephrons (denoted by *SNGFR_sup_* and *SNGFR_jux_* respectively) so that

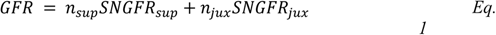

where *n_sup_* = 24,000 and *n_jux_* = 12,000 denote the total number of superficial and juxtamedullary nephrons in a rat kidney. In our previously developed pregnancy-specific superficial nephron models (see Ref. (27)), *SNGFR_sup_* was increased by 30% in MP and 20% in LP based on the experimental results presented in Ref. (32). Then using *Eq. 1* we can note that to have a 30% and 20% increase in the MP and LP GFR, respectively, that *SNGFR_jux_* must also be increased by 30% and 20% in the MP and LP models, respectively (see Table 1). We note that it has been shown that increased GFR in pregnancy has been attributed to increased renal blood flow (3,31,33), so then a similar increase in filtration in both the superficial and juxtamedullary nephrons would be supported by this hypothesis. Note that the total number of nephrons of each type (i.e., *n_sup_* and *n_jux_*) does not change during pregnancy. This is because you are born with the totality of your nephrons, which can be lost through disease, but only gained by transplantation of renal mass.

For changes in morphology and transporter activities we follow the same approach as in our previous study (Ref. (27)). These changes are described in the *Supplementary Materials* and are listed in Table 1. Because adaptations in pregnancy are driven primarily by hormonal changes, we assumed that the pregnancy-induced adaptations in morphology and transporter activities in the juxtamedullary nephrons are the same as the superficial nephrons.

### Simulating transporter inhibition and knockout

We conducted ENaC and H^+^-K^+^-ATPase inhibition *in silico* experiments using our baseline NP, MP, and LP models. In the ENaC inhibition experiment we inhibited the ENaC by 70% and also full knockout (i.e., inhibit by 100%) based on the chronic ENaC blockade experimental processes conducted in West et al. (41). These experiments considered a systemic ENaC blockade, which blocks an estimated 70% of ENaC transport as well as a full ENaC silencing via gene targeting (i.e., ENaC knockout) (41).

We conducted two types of H^+^-K^+^-ATPase *in silico* experiments. In the first simulation type, we fully knocked out the H^+^-K^+^-ATPase transporter only (referred to as HKA-KO; 100% inhibition) for each of the virgin, MP, and LP models. In the second experiment (referred to as “HKA-KO-preg”) we also added changes to the MP and LP H^+^-K^+^-ATPase knockout experiments based on observations from Walter et al. (40). These changes are summarized in Table 2 and are described below.

**Table 2.** Parameter changes for the pregnancy-specific H^+^-K^+^-ATPase knockout simulations (HKAKO-preg). Parameter value ratios for the baseline MP-to-virgin and LP-to-virgin models are the same as those listed in Table 1. Relevant transporters are listed here with the MP-to-virgin and LP-to-virgin ratios for the HKAKO-preg simulations. These changes are based on the results from Ref. (40) and are described in the Materials and Methods section. All other pregnancy-specific parameters are kept the same. NCC: Na^+^-Cl^-^ cotransporter; ENaC: epithelial Na^+^ channel.

|  | Ratios |  |  |  |
| --- | --- | --- | --- | --- |
|  | Baseline<br>MP-to-virgin | HKAKO-preg<br>MP-to-virgin | Baseline<br>LP-to-virgin | HKAKO-preg<br>LP-to-virgin |
| **NCC activity** | 1.00 | 1.00 | 0.45 | 1.00 |
| **Pendrin activity** | 1.65 | 1.00 | 1.70 | 1.00 |
| **ENaC activity** | 2.08 | 1.54 | 2.15 | 1.58 |
| **$H^+-K^+$-ATPase activity** | 2.25 | 0.0 | 2.75 | 0.0 |

Walter et al. (40) noted that during MP and LP, pendrin expression is increased in wild-type (WT) mice. This is also captured in our MP and LP models by increasing pendrin expression (see Table 1). However, in the pregnant H^+^-K^+^-ATPase type 2 knockout (HKA2KO) mice, there was no pregnancy-induced increased pendrin expression from virgin control values (40). Additionally, while normally in MP NCC is unchanged, during LP, NCC is about half of virgin values (40,47). However, expression of NCC in pregnant HKA2KO mice was not found to decrease in the same way as pregnant WT mice (40). We captured altered pendrin and NCC activity in the HKA-KO-preg simulations by setting pendrin and NCC expression in the MP and LP models to virgin values (see Table 2). Lastly, the ENaC has been shown to be highly upregulated during pregnancy (40,41,48). To capture this in our baseline MP and LP models, the parameter for ENaC activity was more than doubled (see Table 1). While Walter et al. (40) did report increased ENaC expression in pregnant HKA2KO mice, the pregnancy-induced increase in ENaC expression was about half of the amount of increase in HKA2KO mice. Thus, we decreased the pregnancy-induced increase in ENaC activity by 50% in both the MP and LP HKAKO-preg simulations (see Table 2).

### Simulating hypertension in female and pregnant rats

To study the impact of hypertension in a female (virgin) rat as well as during pregnancy we developed models that captured renal changes during hypertension. We note that during pregnancy, there are multiple types of hypertensive disorders of pregnancy. In this study we simulate what happens when a female rat that is hypertensive (pre-gestation) gets pregnant. Specifically, we simulate the hypertensive mid-pregnant rat as the hypertensive virgin rat model with then adding the mid-pregnancy changes listed in Table 1. We focus on mid-pregnant hypertension since there have not been any studies to date that investigated nephron function in late pregnant hypertensive rats.

The parameter changes made to simulate hypertension were based largely on the hypertensive female (virgin) rat results in Veiras et al. (37), where the effects of angiotensin II-induced hypertension (a common protocol used in experimental models to induce hypertension (35–38)) on female and male rats was investigated. Additionally, we made some parameter changes using the results from Abreu et al. (49), who investigated renal Na^+^ and aquaporin (AQP) transporters in hypertensive virgin and mid-pregnant rats. The exact parameter changes are listed in Table 3 and are described here. Parameters not listed were unchanged from the baseline model values.

**Table 3.** Parameter changes made in the hypertensive virgin (female) and hypertensive mid-pregnant models given as ratios from the hypertensive (HT) model parameter to the normotensive (NT) model parameter. Parameter change ratios for hypertension are the same in both the hypertensive non-pregnant and hypertensive mid-pregnant models. PCT, proximal convoluted tubule; S3, proximal straight tubule; mTAL, medullary thick ascending limb; cTAL, cortical thick ascending limb; DCT, distal convoluted tubule; CNT, connecting tubule; CCD, cortical collecting duct; OMCD, outer medullary collecting duct; IMCD, inner medullary collecting duct; P_f_, water permeability; P_K_, K^+^ permeability; NHE3, Na^+^/H^+^ exchanger isoform 3; ENaC, epithelial Na^+^ channel; NKCC2, Na^+^-K^+^-2Cl^-^ cotransporter 2; NaPi2, Na^+^-H_2_PO_4_^-^ cotransporter 2; NCC, Na^+^-Cl^-^ cotransporter; KCC, K^+^-Cl^-^ cotransporter.

|  | Ratio: HT-to-NT |
| --- | --- |
| **PCT** |  |
| NHE3 activity | 0.82 |
| NaPi2 activity | 0.85 |
| $Na^+-K^+$-ATPase activity | 1.00 |
| **S3** |  |
| NHE3 activity | 0.82 |
| NaPi2 activity | 0.85 |
| $Na^+-K^+$-ATPase activity | 1.00 |
| **mTAL** |  |
| NKCC2 activity | 0.70 |
| NHE3 activity | 0.82 |
| $Na^+-K^+$-ATPase activity | 0.70 |
| **cTAL** |  |
| NKCC2 activity | 1.74 |
| NHE3 activity | 1.00 |
| $Na^+-K^+$-ATPase activity | 1.00 |
| **DCT** |  |
| $P_K$ (apical, DCT2 only) | 0.30 |
| $Na^+-K^+$-ATPase activity | 1.00 |
| NCC activity | 1.95 |
| ENaC activity | 1.45 |
| **CNT** |  |
| $P_K$ (apical) | 0.30 |
| $Na^+-K^+$-ATPase activity | 1.00 |
| ENaC activity | 1.45 |
| **CCD** |  |
| $P_f$ (transcellular) | 1.30 |
| $Na^+-K^+$-ATPase activity | 1.00 |
| ENaC activity | 1.45 |
| **OMCD** |  |
| $P_f$ (transcellular) | 1.30 |
| Na<sup>+</sup>-K<sup>+</sup>-ATPase activity | 1.00 |
| ENaC activity | 1.45 |
| **IMCD** |  |
| P<sub>f</sub> (transcellular) | 1.30 |
| Na<sup>+</sup>-K<sup>+</sup>-ATPase activity | 1.00 |
| **Hemodynamics** |  |
| SNGFR | 1.00 |

### Proximal segment changes

The fluid inflow at the start of the PT is assumed to be the same as the normotensive virgin and MP models in the hypertensive virgin (virgin-HTN) and hypertensive MP (MP-HTN) models since there are no significant changes in GFR in rats with hypertension (35,36). We note that this would be assuming a similar plasma volume status in chronic hypertension with pregnancy since the pregnancy-induced increase in SNGFR has been shown to be induced by increased renal plasma flow.

Located on the basolateral side of every cell type along the nephron, the Na^+^-K^+^ pump, Na^+^-K^+^-ATPase, is a key driver of Na^+^ and K^+^ transport. Na^+^-K^+^-ATPase abundance has been shown to be decreased in the medullary thick ascending limb by around 30% during hypertension, while abundance in the proximal and distal tubules, as well as the cortical part of the ascending limb was largely unchanged (35,37). To incorporate this into the model, we decreased Na^+^-K^+^-ATPase activity in the medullary thick ascending limb by 30% and left the activity unchanged in the other segments in the hypertensive models (see Table 3).

The Na^+^/H^+^ exchanger isoform 3 (NHE3) is located along the apical side of the epithelial cells in the PT and thick ascending limb. This transporter plays a major role in Na^+^ reabsorption along these early segments of the nephrons. During hypertension, there is a blunted pressure-natriuresis response which often is attributed to lower NHE3 activity (50,51). Veiras et al. (37) found that along the proximal tubule and the medullary thick ascending limb, NHE3 activity in hypertensive rats was decreased from normotensive NHE3 activity. Based on these experimental results, we decreased NHE3 activity by 15% in the proximal tubule and medullary thick ascending limb for the hypertensive virgin (female) rats. We did not change NHE3 activity in the cortical segment of the thick ascending limb since NHE3 activity has been reported to be largely unchanged in this segment in hypertensive rats (35).

The Na^+^/P_i_ cotransporter 2 (NaPi2) is expressed along the proximal tubule. Veiras et al. (37) found that NaPi2 expression was decreased in hypertensive rats compared to normotensive rats. Based on these experimental results, we decreased NaPi2 activity in hypertensive rats by 15%, which is within the range of NaPi2 expression changes in hypertensive female rats reported (37).

The Na^+^-K^+^-2Cl^-^ cotransporter (NKCC2) is located on apical membrane along the thick ascending limb. During hypertension, NKCC2 activity changes are region-specific (35–37).

Specifically, in the cortex, NKCC2 activity is increased, while in the medulla, NKCC2 is significantly decreased (35–37). We decreased NKCC2 activity by 30% in the medullary thick ascending limb and increased NKCC2 activity by 74% in the cortical thick ascending limb. These changes are within range of reported experimental values (35–37).

### Distal segment changes

The Na^+^-Cl^-^ cotransporter (NCC) is expressed along the distal convoluted tubule. During hypertension, NCC abundance and phosphorylation has been found to increase significantly (35–37). We nearly doubled NCC activity to capture these findings (see Table 3).

Abreu et al. (49) found a major decrease in the expression of the renal outer medullary K^+^ channel (ROMK) during hypertension for both virgin and MP. To capture this, we decreased K^+^ permeability in the late distal convoluted tubule and the connecting tubule by 70% (see Table 3). During hypertension, ENaC activity is increased (35–37). The increase in ENaC expression during hypertension is higher in females than in males (37). We increased ENaC activity by 45% in the distal segments and collecting duct which is within range of reported experimental findings (37).

Abreu et al. (49) also reported increased aquaporin 2 (AQP2) expression in the collecting duct. Specifically, in the hypertensive virgin, the AQP2 expression increased from normotensive virgin levels by around 30%. We changed the water permeability in the collecting duct segments based on this result.

## Research Ethics

There was no use of animal or human subjects in this study.

## Data Availability

The code for the nephron model used in this study is available at the GitHub repository https://github.com/Layton-Lab/nephron. We have also used Zenodo to assign a DOI to the repository: https://doi.org/10.5281/zenodo.7888417 which includes exact code version used as well as simulation data with scripts to generate the figures presented in the *Results*.

## Results

### Baseline mid-pregnant and late pregnant models predict a balance of the marked increase in filtered loads with pregnancy-induced enhancement in transport capacity

Solute and fluid volume reabsorption along specific nephron segments depends on activity of membrane transporters, permeability of the apical and basolateral membrane, as well as tubular morphology. Many changes occur in almost all nephron segments during pregnancy. To investigate the impact of pregnancy-induced changes on renal transport, we developed baseline models to represent kidney function in a virgin, MP, and LP rat.

In the MP and LP models, the filtration to the nephrons is 30% and 20% higher than the virgin model values, respectively (Table 1). This yields a proportional increase in total fluid and solute delivery at the start of the proximal tubule (PT) in the MP and LP model predictions (Figure 2). Model predictions indicate that, due to tubular hypertrophy (i.e., increased transport area) as well as upregulation in the Na^+^/H^+^ exchanger (NHE3) and the K^+^-Cl^-^ cotransporter (KCC), predicted net Na^+^ reabsorption in the PT of the MP and LP models is significantly larger than in the virgin rat model (Figure 3A; see Ref. (27)). Specifically, MP and LP models predicted a 27% and 16% increase in net Na^+^ reabsorption along the PT when compared to virgin (Figure 3A). The increase in Na^+^ reabsorption is about the same proportion increase as the in GFR in both the MP and LP models, hence glomerulotubular balance in the PT appears to hold in pregnancy, with a fractional reabsorption of Na^+^ about 60% for each of the virgin, MP, and LP models along the PT. Analogous results are obtained for Cl^-^ and fluid volume along the PT (see Figure 2; Figure 3).

**Figure 2.**
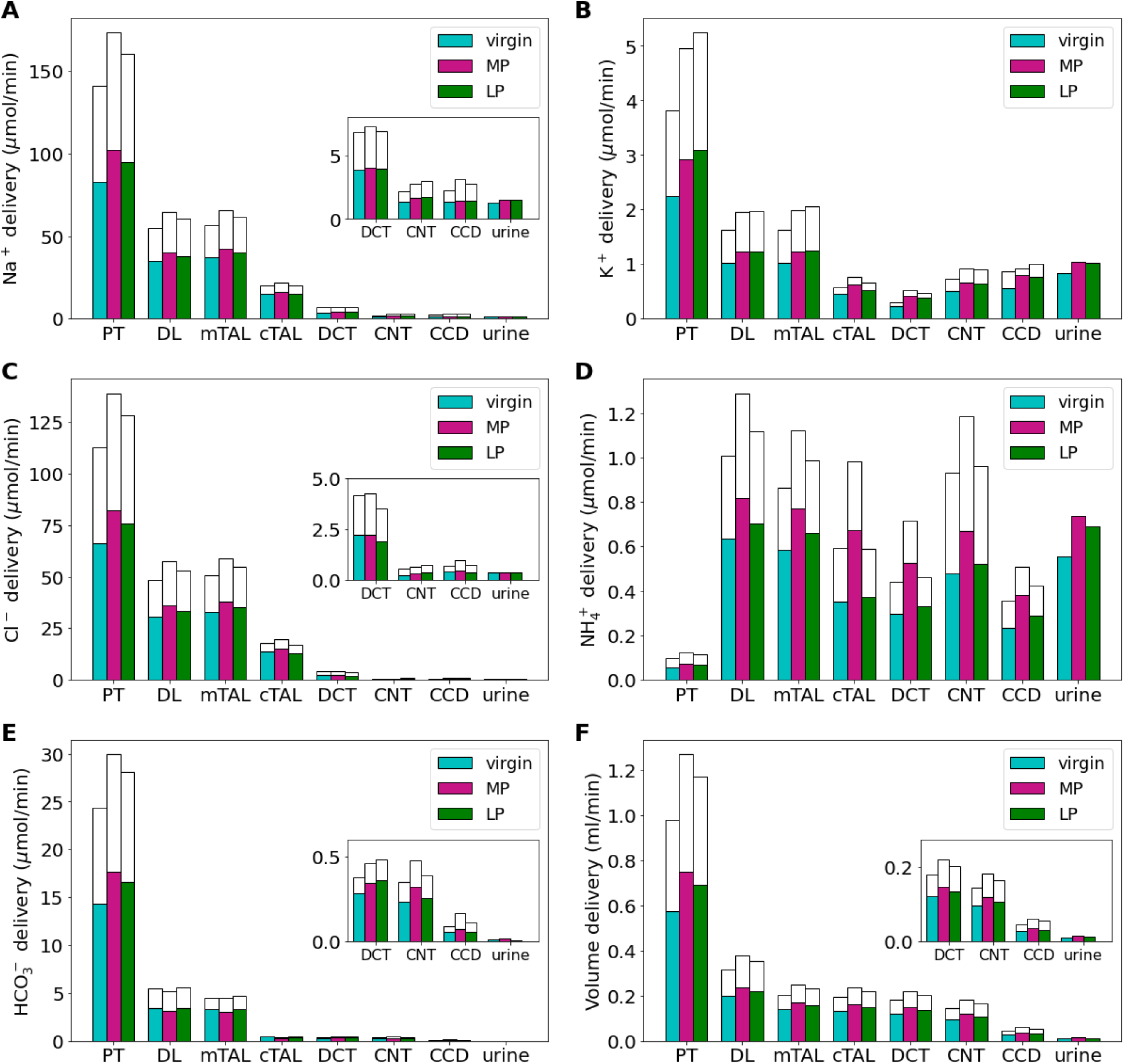
Delivery of key solutes (A-E) and fluid volume (F) to the beginning of individual nephron segments in the virgin, mid-pregnant (MP), and late pregnant (LP) rat models. The colored bars denote the superficial nephron values, and the white bars denote the weighted totals of the juxtamedullary nephrons (five representative model nephrons) where total bar height is the sum of the delivery to superficial and juxtamedullary nephrons. Note that since the model assumes that the superficial-to-juxtamedullary nephron ratio is 2:1, the superficial delivery values are generally higher. Additionally, the juxtamedullary and superficial nephrons merge at the collecting duct so the urine output is combined. PT, proximal tubule; DL, descending limb; LAL, thin ascending limb; mTAL, medullary thick ascending limb; DCT, distal convoluted tubule; CNT, connecting tubule; CCD, cortical collecting duct. Insets: reproduction of distal segment values.

Downstream of the PT, the thick ascending limb reabsorbs much of the Na^+^ and Cl^-^ remaining in the lumen. In the MP model, Na^+^ delivery to the thick ascending limb is 16% higher than virgin model values due to the increased filtered load of Na^+^ along with similar fractional reabsorption of Na^+^ by the PT (Figure 2A). Reabsorption along the thick ascending limb increases by about 16% in MP (Figure 3A). Analogous results are obtained for Cl^-^. In the end, urinary Na^+^ excretion for both MP and LP is about 16% higher than virgin values and urinary Cl^-^ excretion is increased by 4% and 6% in the MP and LP models, respectively. The increase in natriuresis and Cl^-^ excretion in the pregnancy models versus the virgin model are within reported ranges (48,52).

**Figure 3.**
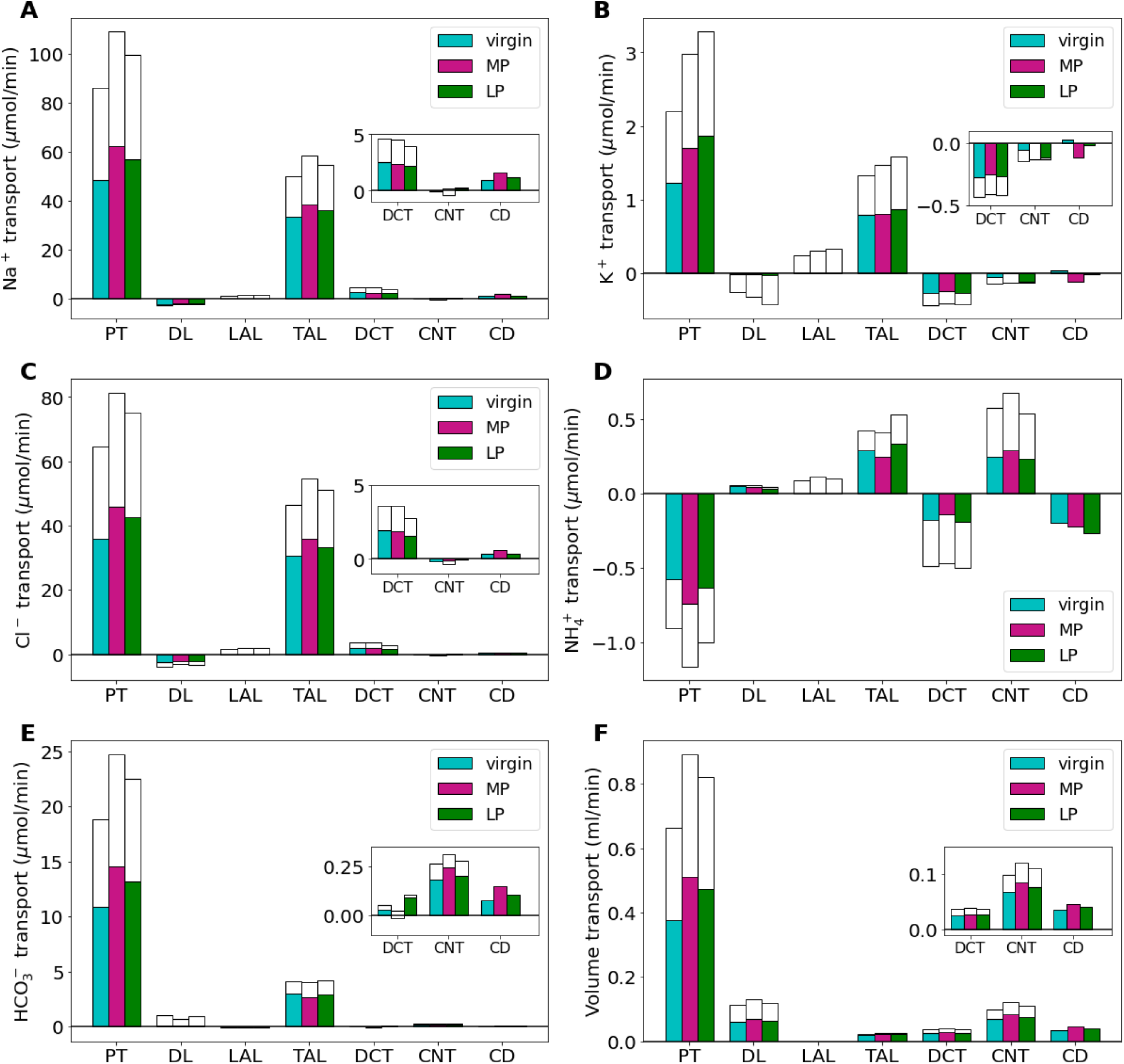
Net segmental transport of key solutes (A-E) and volume (F) along the individual nephron segments in the virgin, MP, and LP models. Transport is positive out of the nephron segment, i.e., positive transport indicates net reabsorption and negative transport indicates net secretion along that segment. Colored bars denote the superficial nephron values and white bars denote juxtamedullary values computed as a weighted sum of the five representative juxtamedullary nephrons. PT, proximal tubule; DL, descending limb; LAL, thin ascending limb; TAL, thick ascending limb; DCT, distal convoluted tubule; CNT, connecting tubule; CD, collecting duct. Insets: reproductions of distal segment values.

Along the PT, the increased Na^+^ reabsorption provides the driving force for increased water reabsorption along the PT by 34% and 24% in the MP and LP models, respectively, when compared to virgin model values (Figure 3F). In the segments after the macula densa and the collecting duct, fluid reabsorption is also increased by 22% and 12% in the MP and LP models, respectively when compared to virgin model fluid reabsorption (Figure 3F). This is likely driven by increased AQP2 expression in the collecting duct, increased nephron diameter and length, as well as increased fluid volume delivery to these segments.

About 58% of the filtered K^+^ is reabsorbed along the PT in the virgin model (Figure 2B). During LP, the fractional reabsorption of K^+^ in the PT increases to about 63% (Figure 3B). This is due to the massive increase in delivery of K^+^ from a combination of 30% increase in plasma K^+^ concentration, increased GFR, and increased K^+^ transport capacity (Table 1). Distal segment K^+^ secretion along the distal convoluted tubules (DCT) and connecting tubules (CNT) is decreased by about 8% in the MP and LP models from virgin model values despite known kaliuretic factors such as increased Na^+^ delivery (Figure 3B). This is due to the decreased K^+^ channel permeability in these segments (see Table 1). In all, K^+^ urinary excretion is predicted to be 2.4, 3.0, and 2.9 mmol/day in the virgin, MP, and LP models, respectively. The increase in kaliuresis from virgin values in the MP and LP models is within reported ranges (28,52).

### Proximal tubule hypertrophy and the upregulation of epithelial Na^+^ channel and H^+^-K^+^-ATPase play key roles in maintaining Na^+^ and K^+^ balance during pregnancy

Using our baseline LP computational model, we conducted “*what-if”* simulations to predict how key individual changes impact renal transport during late pregnancy. To do this, we considered the renal adaptations that were found to have the largest impact on superficial nephron transport in our previous study (see Ref. (27)) and conducted simulations of LP kidney function without such adaptation (i.e., respective parameter value at the virgin model value) using our LP multi-nephron computational model. Transport of Na^+^ and K^+^ for adaptations that affect the proximal segments of the nephron (i.e., PT length and NHE3 activity) are shown in Figure 4 and adaptations that affect only the distal segments (i.e., ENaC activity, K^+^ channel permeability, H^+^-K^+^-ATPase activity) are shown in Figure 5.

**Figure 4.**
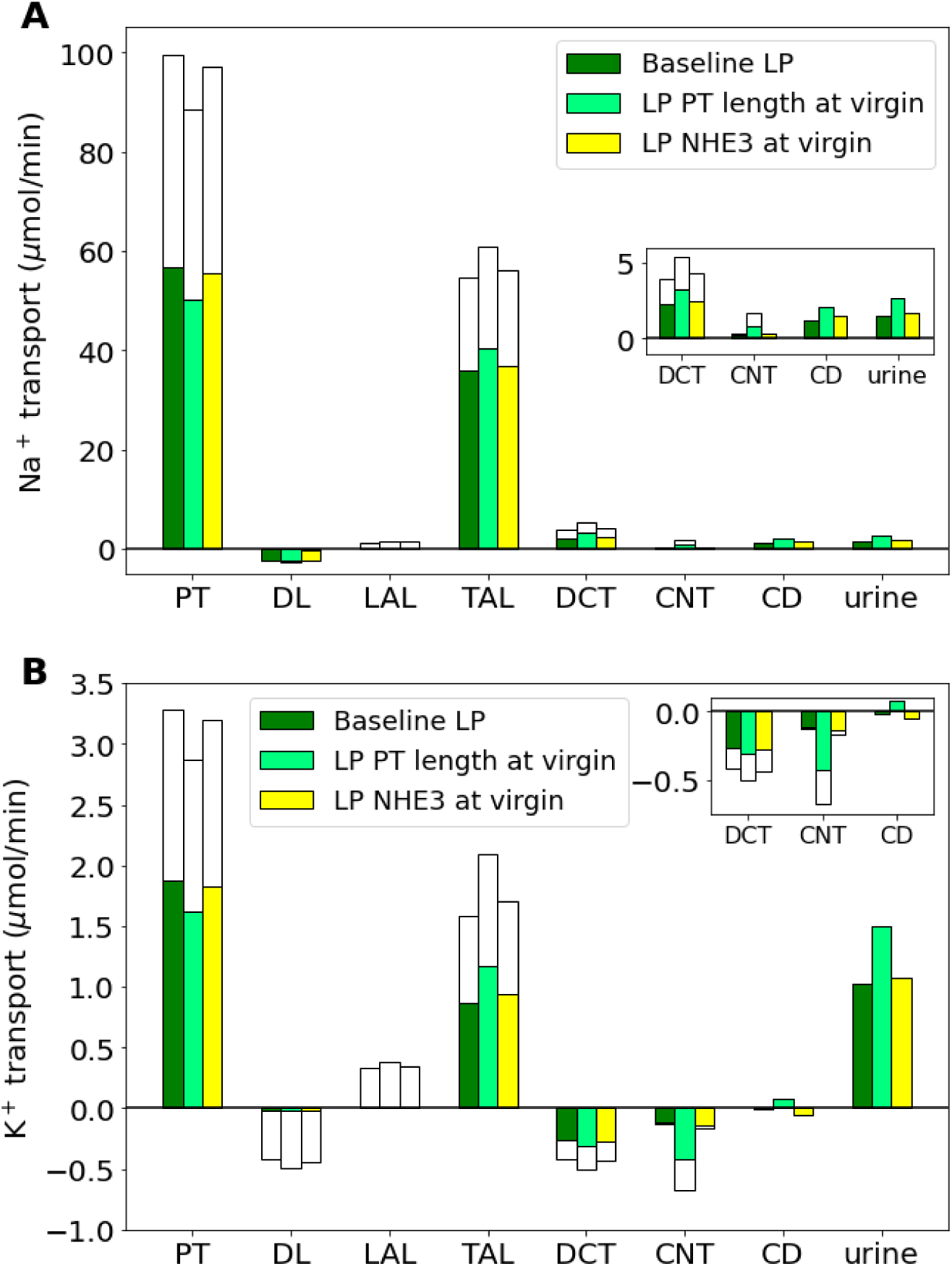
Impact of key pregnancy-induced proximal segment renal adaptations on net segmental transport and urine excretion of Na^+^ (A) and K^+^ (B) along individual nephron segments in the baseline LP model and LP model with one pregnancy-induced change (as labeled) set to the respective virgin parameter value: NHE3 activity and PT length. Transport is positive out of the nephron segment, i.e., positive transport shows net reabsorption and negative net secretion along that segment. PT, proximal tubule; DL, descending limb; LAL, thin ascending limb; mTAL, medullary thick ascending limb; DCT, distal convoluted tubule; CNT, connecting tubule; CCD, cortical collecting duct; NHE3, Na^+^/H^+^ exchanger isoform 3; NKATPase, Na^+^-K^+^-ATPase.

Most reabsorption in the kidneys along the proximal segments, which include the PT and the loop of Henle. During LP, the PT length is increased by about 17% from the baseline virgin model nephron length (Table 1; Refs. (27,30,34)), which increases transport capacity along the PT of the nephrons. Without this change, we predict that the PT reabsorbs 11% less Na^+^ than the baseline LP model (Figure 4A). While reabsorption along the thick ascending limbs and distal segments is increased, it is not enough to make up for the loss resulting in an increase of about 77% in urine excretion of Na^+^ (Figure 4A). Similarly, without the PT length increase, K^+^ reabsorption is significantly decreased in the PT segment, ultimately resulting in a 46% increase in urinary K^+^ excretion (Figure 4B). The Na^+^/H^+^ exchanger along the proximal tubule drives much of the Na^+^ reabsorption along the proximal segments. In our MP and LP models, NHE3 activity is increased by about 40% and 20%, respectively, from virgin values. Decreasing NHE3 activity to the virgin value results in a 2% decrease in PT reabsorption. This loss of reabsorption is not made up for in the later segments and results in a 17% and 8% increase in urinary Na^+^ and K^+^ excretion, respectively (Figure 4).

The distal segments account for much of the final fine tuning of fluid concentrations in the kidney nephrons before what is remaining in the lumen is excreted as urine. Expression of the ENaC is abundantly increased during pregnancy, likely contributing to increased ENaC activity. Additionally, activity of the H^+^-K^+^-ATPase transporter is significantly increased, while distal segment K^+^ secretory channels, the renal outer medullary K^+^ channel (ROMK) and big K^+^ channel (BK), are significantly decreased (Note that this is represented by P_K_ in Table 1). We conducted simulations of the LP model without such changes and present the results for Na^+^ and K^+^ transport as well as urinary excretion in Figure 5.

**Figure 5.**
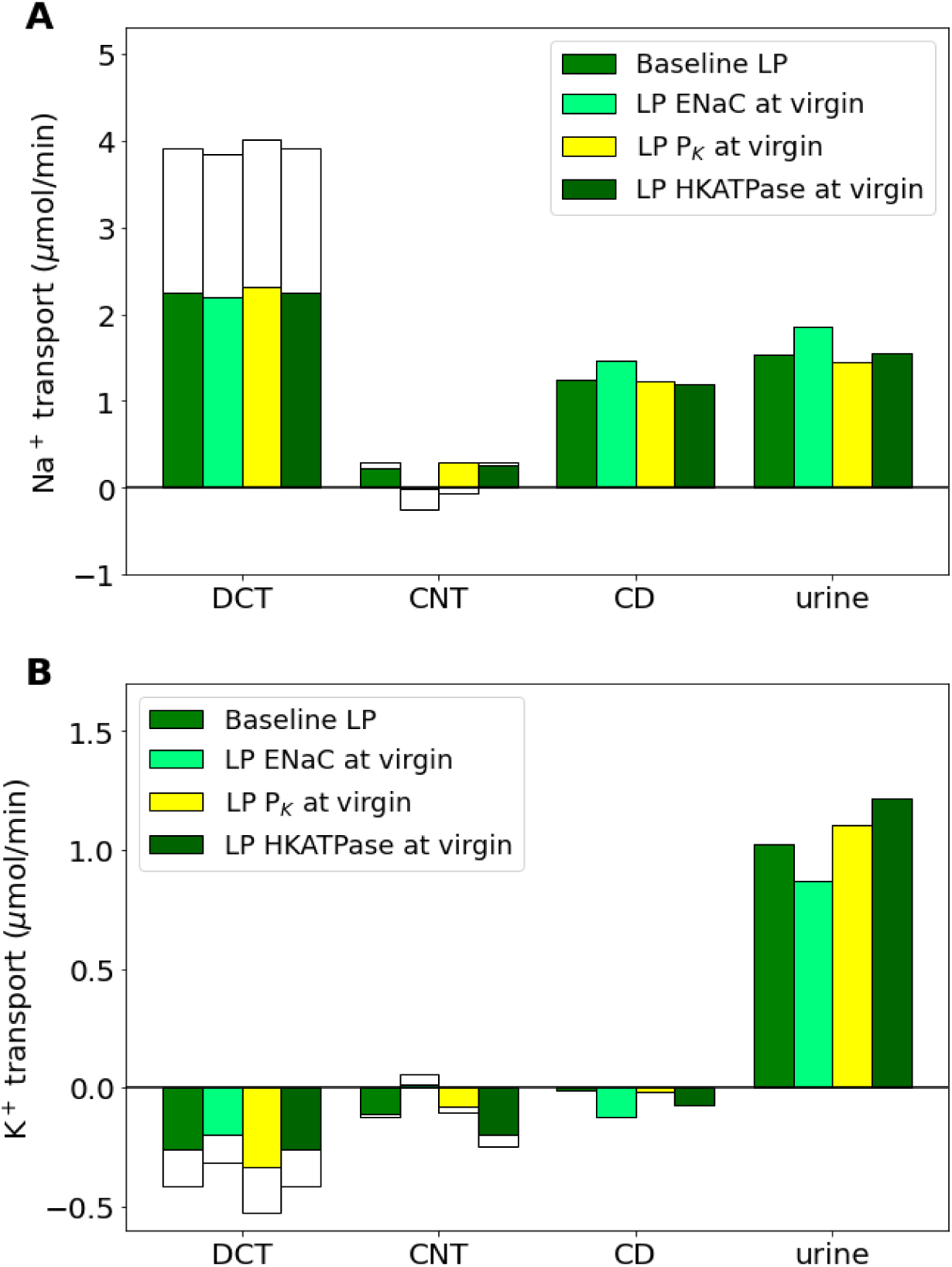
Impact of key pregnancy-induced adaptations in the distal segments on net segmental transport and urine excretion of Na^+^ (A) and K^+^ (B) along individual nephron segments in the baseline LP model and LP model with one pregnancy-induced change set to the respective virgin value: ENaC activity, P_K_, and H^+^-K^+^-ATPase activity. Transport is positive out of the nephron segment, i.e., positive transport shows net reabsorption and negative net secretion along that segment. Only the distal segment results are shown because these changes only affect distal segments. HKATPase, H^+^-K^+^-ATPase; ENaC, epithelial Na^+^ channel; P_K_, permeability of K^+^; DCT, distal convoluted tubule; CNT, connecting tubule; CCD, cortical collecting duct.

Without the large increase in ENaC activity in the LP model, Na^+^ reabsorption in the DCT and CNT decreased by 13%. This yielded a 22% increase in urinary Na^+^ excretion (Figure 5A). Notably, increased ENaC activity is a known kaliuretic factor. Without a pregnancy-induced increase in ENaC activity we predict a 15% decrease in urinary K^+^ excretion (Figure 5B).

Without the pregnancy-induced massive decrease in K^+^ secretory channel abundance, K^+^ secretion along the DCT increased by 26%. Decreased CNT K^+^ secretion makes up for some of this extra K^+^ loss, but model simulations predict an increase in K^+^ excretion by about 8%. The other K^+^ transporter altered during LP is H^+^-K^+^-ATPase activity, which is more than double virgin values (Table 1). Without this change, both the distal tubule and collecting duct (CD) K^+^ secretion more than doubles resulting in urinary K^+^ increase by 19% (Figure 5B).

### Increased epithelial Na^+^ channel activity is essential for sufficient Na^+^ reabsorption during pregnancy

West et al. (41) investigated the effects of inhibition of the ENaC on pregnancy and found that a chronic ENaC blockade ablated the Na^+^ retention found in normal pregnancy. We conducted simulations to predict the functional implications of a 70% ENaC inhibition and a full ENaC knockout (i.e., 100% inhibition) on Na^+^ and K^+^ transport along the distal segments. Results are shown in Figure 6.

**Figure 6.**
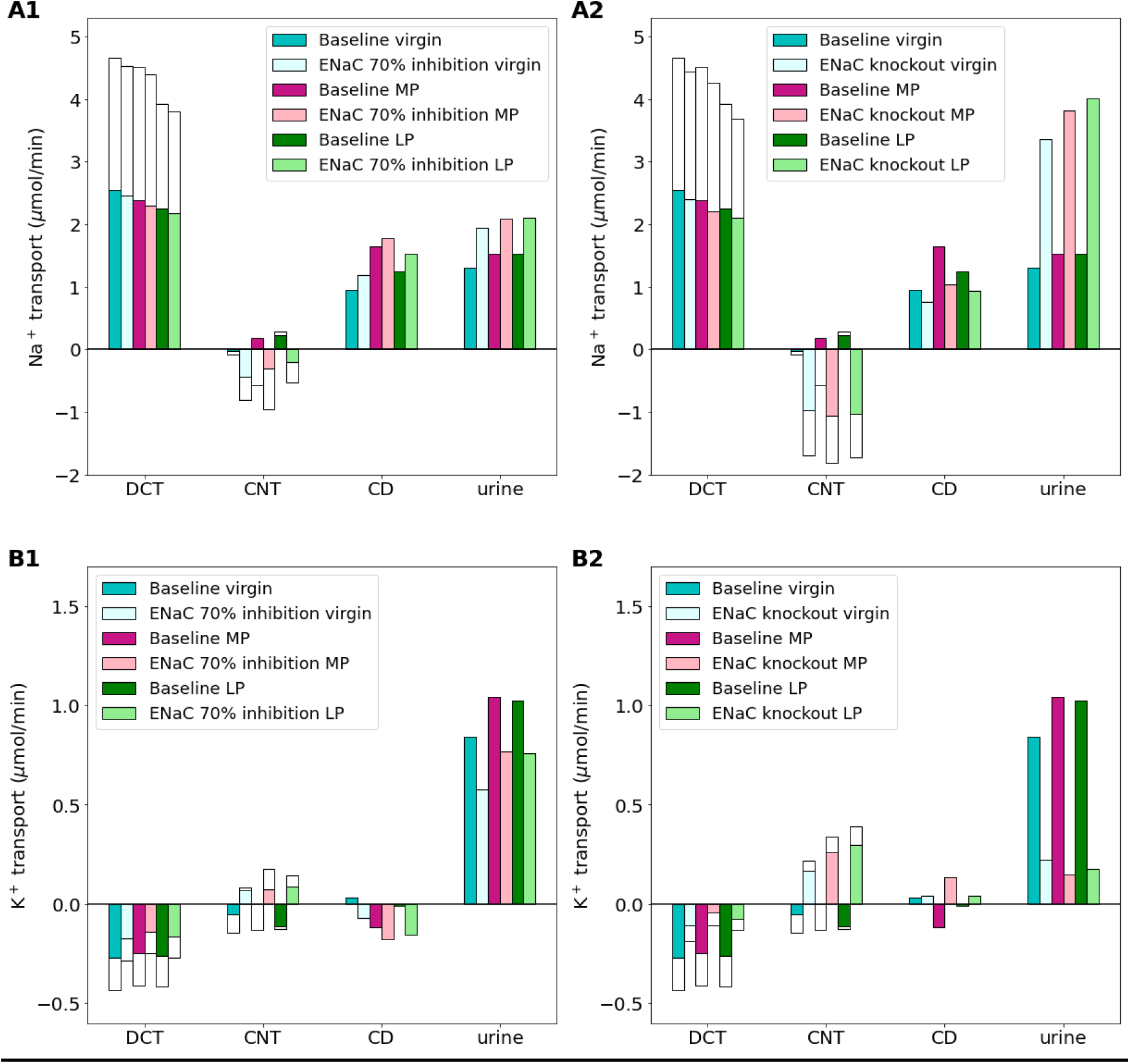
Impact of epithelial Na^+^ channel (ENaC) inhibition (70%) (A1-B1) and ENaC knockout (A2-B2) on distal segment net segmental transport of Na^+^ (A1-A2) and K^+^ (B1-B2) along individual nephron segments in the virgin, mid-pregnant (MP), and late pregnant (LP) models. Transport is positive out of the nephron segment, i.e., positive transport shows net reabsorption and negative net secretion along that segment. The last bar shows delivery of urine which is the same as in Figure 2. Only the distal segments are shown because ENaC inhibition only affects distal segments. DCT, distal convoluted tubule; CNT, connecting tubule; CCD, cortical collecting duct.

When the ENaC is inhibited by 70% in both the MP and LP models, total Na^+^ reabsorption along the DCT and CNT decreases by about 20%, when compared to the respective baseline MP and LP models. Collecting duct Na^+^ reabsorption does increase in both models, but not enough to make up for the lost reabsorption resulting in a 48% and 38% increase in urinary Na^+^ excretion in the MP and LP models, respectively (Figure 6A1). ENaC inhibition has an opposite effect on K^+^. Specifically, K^+^ secretion is decreased by ENaC inhibition so much that the models predict a net K^+^ reabsorption along the CNT for each of the virgin, MP, and LP models (Figure 6B1). This significant change results in a net reduction in K^+^ secretion along the distal segments of 82% in the MP model. In the end urinary excretion of K^+^ is reduced by about 26% when compared to the baseline MP and LP models (Figure 6B1).

A full ENaC knockout leads to even more drastic effects on Na^+^ and K^+^ handling in each of the virgin, MP, and LP model predictions when compared to 70% inhibition. Specifically, the CNT has net secretion of Na^+^ with net reabsorption of K^+^. This happens likely because the ENaC is the primary Na^+^ transporter along this segment. As a result, urinary Na^+^ excretion is predicted to more than double while urinary K^+^ excretion is about 25% of baseline for the virgin, MP, and LP models (Figure 6A2; Figure 6B2).

### Adaptations in pregnant H^+^-K^+^-ATPase knockout animals decreases excess K^+^ loss while increasing Na^+^ excretion

Walter et al. (40) investigated the affect of H^+^-K^+^-ATPase knockout (HKA-KO) on Na^+^ and K^+^ regulation in pregnant mice and found that pregnant HKA-KO mice had only a modestly expanded plasma volume and altered K^+^ balance when compared to pregnant wild-type mice.

We conducted two types of HKA-KO simulations. In the first simulation, we did a full 100% inhibition of the H^+^-K^+^-ATPase transporter only. We refer to this simulation as HKA-KO. Results are shown in Figure 7. In the second simulation, we also added a changes in Na^+^-Cl^-^ cotransporter, ENaC, and Pendrin transporter activities in pregnant H^+^-K^+^-ATPase knockout mice as reported in Walter et al. (40) (see *Materials and Methods*; Table 2). We refer to this simulation as “HKA-KO-preg”. These results are shown in Figure 7.

**Figure 7.**
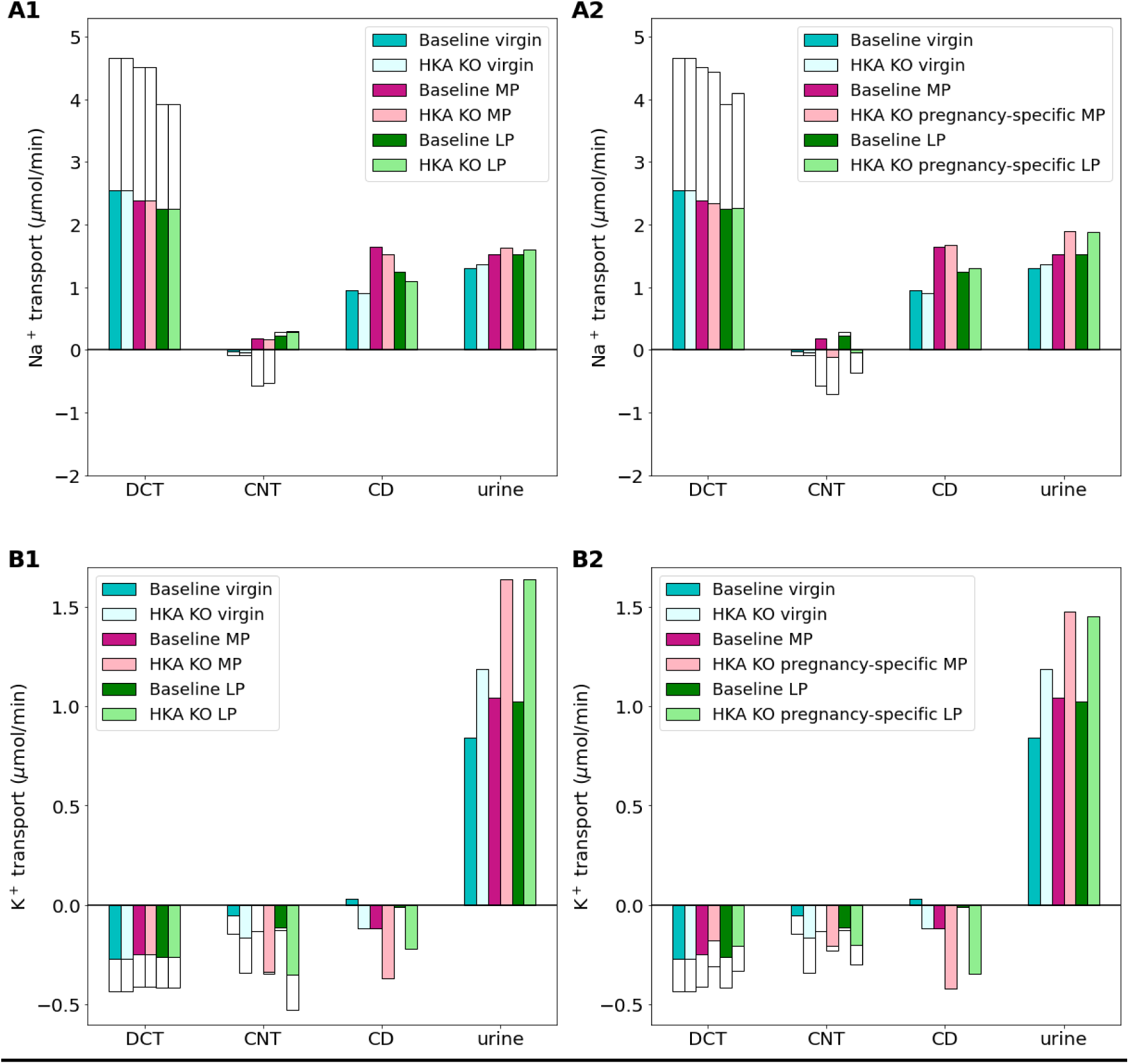
Impact of H^+^-K^+^-ATPase knockout (i.e.,HKA KO) (A1-B1) and pregnancy-specific HKA KO (i.e., HKAKO-preg) (A2-B2) on distal segment net segmental transport of Na^+^ (A1-A2) and K^+^ (B1-B2) along individual nephron segments in the virgin, mid-pregnant (MP), and late pregnant (LP) models. Transport is positive out of the nephron segment, i.e., positive transport shows net reabsorption and negative net secretion along that segment. Only the distal segments are shown because HKA inhibition only affects distal segments. DCT, distal convoluted tubule; CNT, connecting tubule; CCD, cortical collecting duct.

Net K^+^ secretion along the DCT and CNT is predicted to increase by 85% and 77% in the MP HKA-KO and LP HKA-KO simulations, respectively, when compared to the baseline MP and LP models. This results in about a 60% increase in urinary K^+^ excretion for both the MP and LP HKA-KO model simulations. The virgin HKA-KO model predicts an increase in K^+^ secretion along these segments, but by a smaller fraction of 34%, yielding a 41% increase in urinary K^+^ excretion when compared to the baseline virgin model (Figure 7). Na^+^ handling is slightly altered resulting in about a 5% increase in urinary Na^+^ excretion for each of the virgin, MP, and LP model simulations.

Walter et al. (40) found that in pregnant H^+^-K^+^-ATPase type 2 knockout mice, pregnancy-induced adaptations in the Na^+^-Cl^-^ cotransporter, ENaC, and Pendrin transporter were altered from normal adaptations. We conducted simulations with these observed changes and called these simulations the “HKA-KO-preg” simulations (see Figure 7). The additional adaptations alter how Na^+^ and K^+^ is handled along the distal segments when compared to the HKA-KO only simulations. Specifically, in the HKA-KO-preg MP simulations, K^+^ secretion along the DCT and CNT is 32% higher than baseline MP simulations. Similarly, the HKA-KO-preg LP simulation predict a 19% increase in K^+^ secretion along these segments when compared to baseline LP. This is likely due to the decreased ENaC activity because increased ENaC activity would increase K^+^ secretion in these segments. In the end urinary K^+^ excretion for the HKA-KO-preg MP and LP simulations is about 42% higher than the respective baseline MP and LP model predictions. Additionally, since there is Na^+^ transporter changes, Na^+^ handling is altered such that urinary Na^+^ excretion is increased by about 24% in both the HKA-KO-preg MP and HKA-KO-preg LP models when compared to the respective baseline MP and LP model predictions.

### Chronic hypertension induces a shift in Na^+^ load to distal segments in virgin and MP rat kidneys

We developed hypertensive virgin and MP models using the changes listed in Table 3 and described in *Materials and Methods*. We denote the hypertensive virgin model by virgin-HTN and the hypertensive MP model by MP-HTN. Delivery of Na^+^, K^+^, and fluid volume to each segment for the baseline virgin, virgin-HTN, baseline MP, and MP-HTN models are shown in Figure 8. Net segmental transport predictions for each of the normotensive and hypertensive virgin and MP models is given in Figure 9.

**Figure 8.**
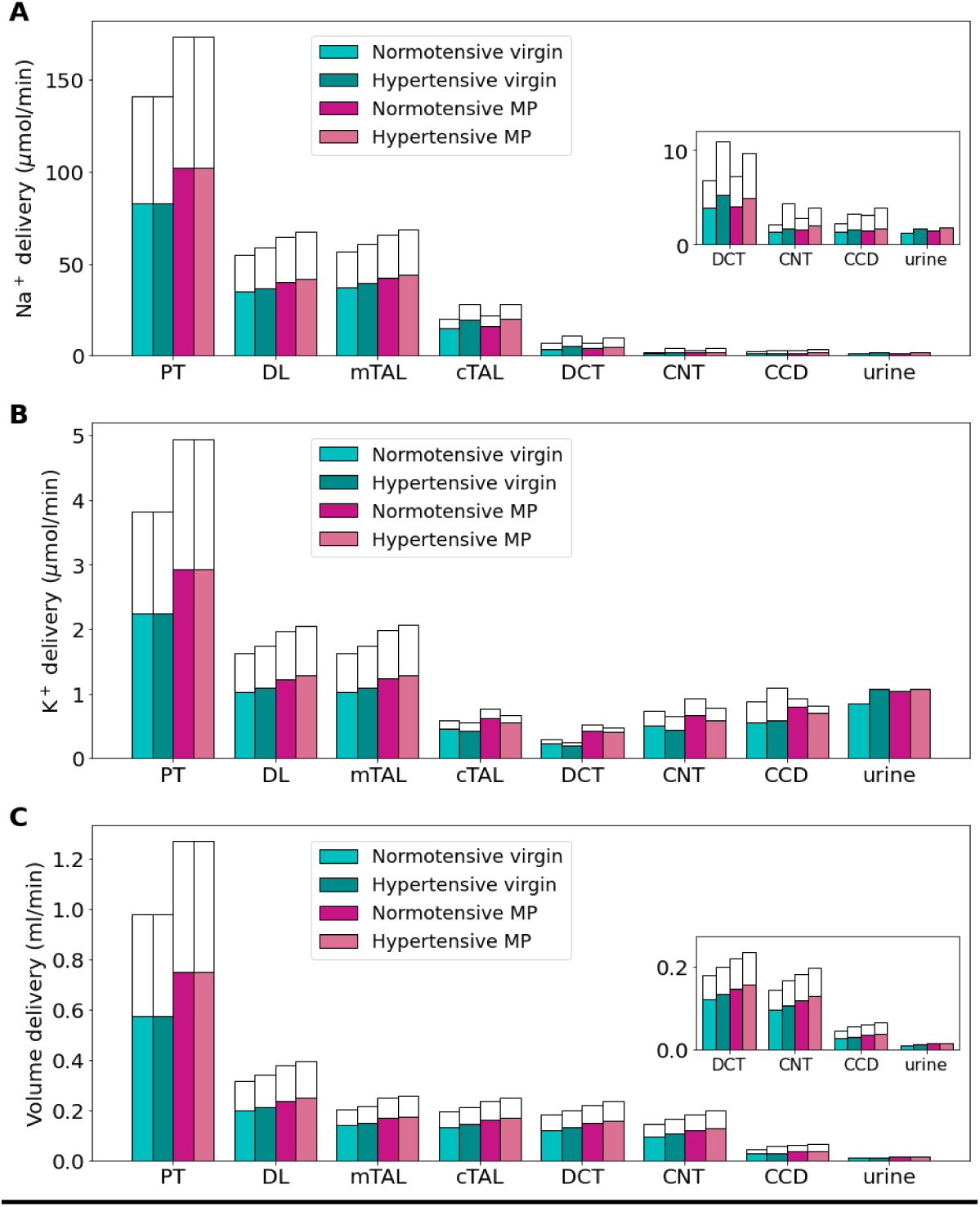
Delivery of key solutes (A-E) and fluid volume (F) in the normotensive and hypertensive virgin and mid-pregnant (MP) models. The colored bars denote the superficial nephron values, and the white bars denote the weighted totals of the juxtamedullary nephrons (five representative model nephrons). PT, proximal tubule; DL, descending limb; LAL, thin ascending limb; mTAL, medullary thick ascending limb; cTAL, cortical thick ascending limb; DCT, distal convoluted tubule; CNT, connecting tubule; CCD, cortical collecting duct. Insets: reproduction of distal segment values.

**Figure 9.**
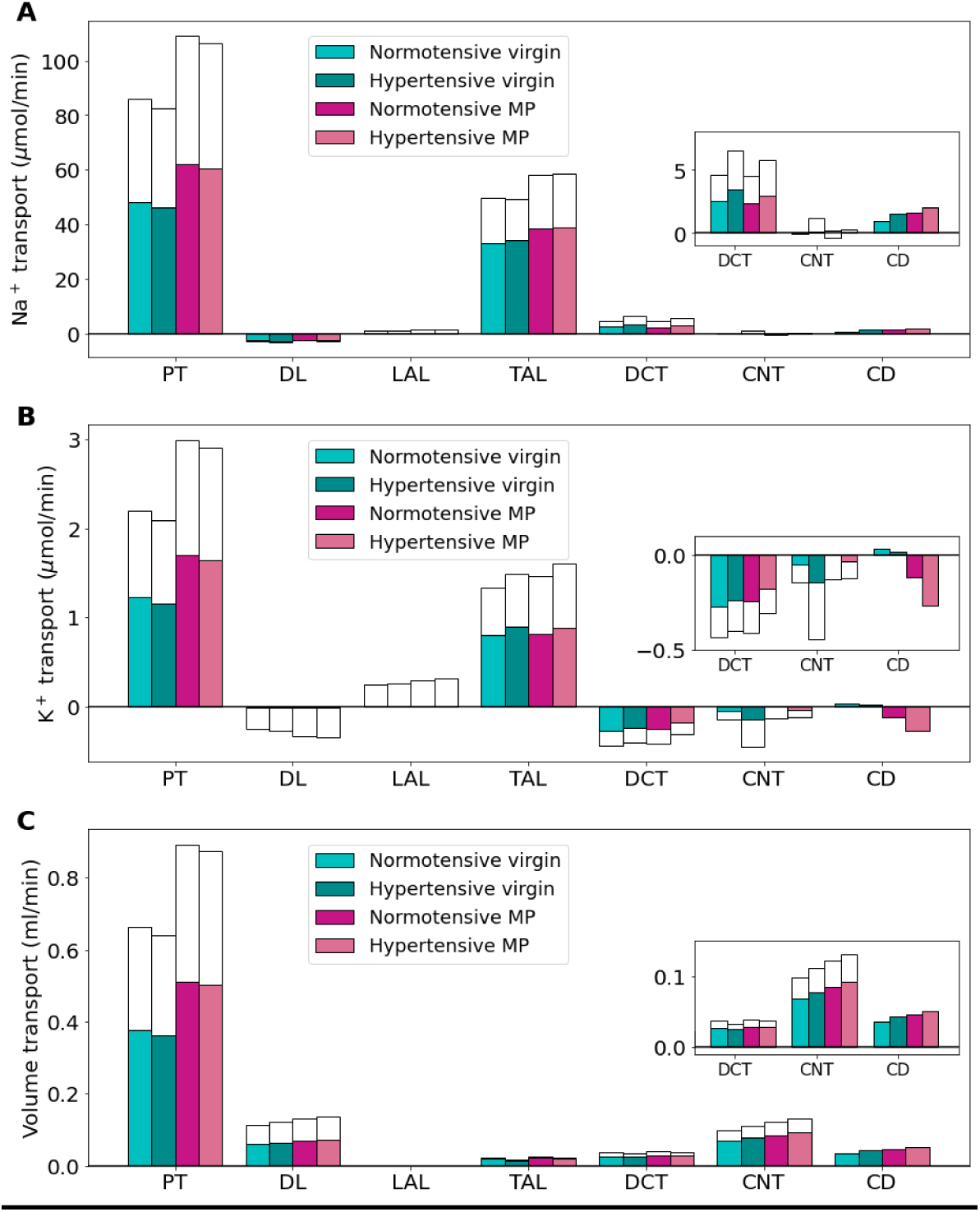
Net segmental transport of key solutes (A-E) and volume (F) along the individual nephron segments in the normotensive and hypertensive virgin and mid-pregnant (MP) models. Transport is positive out of the nephron segment, i.e., positive transport shows net reabsorption and negative net secretion along that segment. Colored bars denote the superficial nephron values and white bars denote juxtamedullary values computed as a weighted sum of the five representative juxtamedullary nephrons. PT, proximal tubule; DL, descending limb; LAL, thin ascending limb; mTAL, medullary thick ascending limb; cTAL, cortical thick ascending limb; DCT, distal convoluted tubule; CNT, connecting tubule; CD, collecting duct. Insets: reproductions of distal segment values.

Due to pressure natriuresis, PT Na^+^ reabsorption is decreased slightly (about 4%) in the virgin-HTN and MP-HTN models from the respective normotensive model predictions (Figure 9A). The virgin-HTN and MP-HTN models predict a 5% and 3% decrease in PT K^+^ reabsorption when compared to the respective normotensive models (Figure 9B). Net fluid volume reabsorption along the PT decreased by about 3% for both the virgin-HTN and MP-HTN models when compared to the respective normotensive baseline models (Figure 9C). The lower fluid volume reabsorption results in increased volume delivery to each of the segments along the nephron. These changes in Na^+^, K^+^, and volume transport along the PT are driven by the decreased NHE3 and NaPi2 activity in the hypertensive models (see Table 3).

Region-specific changes in the Na^+^-K^+^-2Cl^-^ cotransporter (NKCC2), NHE3, and Na^+^-K^+^-ATPase activity (Table 3). lead to differential alterations in transport in the medullary and cortical segments of the thick ascending limb. In the medullary thick ascending limb of both hypertensive models, NKCC2, NHE3, and Na^+^-K^+^-ATPase activity are assumed lowered decreased (Table 3). Consequently, Na^+^ reabsorption along the medullary thick ascending limb decreases by 12% in the virgin-HTN model and 8% in the MP-HTN model relative to the respective normotensive models (Figure 9A). As such, more Na^+^ reaches the cortical thick ascending limb where, unlike the medullary thick ascending limb, transport capacity is enhanced: NKCC2 activity is increased from normotensive values in the hypertensive models, while NHE3 and Na^+^-K^+^-ATPase activity is unchanged from normotensive values (Table 3). This results in about a 27% increase in Na^+^ reabsorption along this segment in both the hypertensive models (Figure 9A).

The reduced PT K^+^ reabsorption leads to a 7% and 5% increase in K^+^ delivery to the thick ascending limb in the virgin-HTN and MP-HTN models, respectively, when compared to the respective normotensive model (Figure 8B). Along the medullary thick ascending limb, K^+^ transport is increased by about 15% in the hypertensive models compared to the respective normotensive models (Figure 9B). This increased transport in the medullary thick ascending limb results in decreased cortical thick ascending limb K^+^ delivery in the hypertensive models compared to the respective normotensive models for both the virgin-HTN and MP-HTN models.

Na^+^ transport along the DCT and CNT exhibits a substantial increase in both hypertensive models (69% and 42% increase, respectively, in the virgin-HTN and MP-HTN above baseline normotensive model predictions), due to elevated delivery as well as the enhanced Na^+^-Cl^-^ cotransporter and ENaC activity (Figure 9A). Major K^+^ secretion occurs along the DCT and CNT. During hypertension, the expression of K^+^ secretory channel ROMK is significantly decreased (Table 3). In the late DCT, the reduced K^+^ secretion from the ROMK leads to 8% and 25% lower K^+^ secretion in the DCT for the virgin-HTN and MP-HTN models, respectively when compared to the respective normotensive virgin and MP models. This is where the virgin-HTN and MP-HTN models seem to differ. Connecting tubule K^+^ secretion is increased in the virgin-HTN model when compared to virgin so that in total the K^+^ secretion along the DCT and CNT is 45% higher than virgin distal tubule K^+^ secretion. This increase is likely driven by the effect of increased Na^+^ flow in the CNT and increased ENaC activity (a known kaliuretic factor). The increased Na^+^ in the CNT creates a favorable electrical potential gradient which in turn increases K^+^ secretion. K^+^ secretion in the MP-HTN model along the DCT and CNT is predicted to be about the same as normotensive MP distal tubule K^+^ secretion. This is likely due to the already significantly decreased ROMK channel permeability in the normotensive MP model (Table 1).

In summary, decreased Na^+^ transport in the proximal segments effects the early K^+^ reabsorption which leads to higher K^+^ flow to the distal segments. Further along the nephron in the distal segments K^+^ secretory channels are reduced to lower K^+^ secretion despite high Na^+^ flow and kaliuretic factors such as increased ENaC activity.

## Discussion

What are the physiological implications of pregnancy-induced renal adaptations? How does knockout of the ENaC and H^+^-K^+^-ATPase transporters impact distal tubule transport and urinary excretion in virgin and pregnant rats? How does altered expression and activity of renal transporters impact kidney function in a female or pregnant rat with hypertension? To answer these questions, we developed and analyzed computational models of kidney function in a female rat during mid- and late pregnancy as well as chronic hypertension. The main findings of this study are:

- Without pregnancy-induced adaptations in proximal tubule length and NHE3 activity, proximal tubule Na^+^ reabsorption during pregnancy is insufficient, resulting in increased urinary Na^+^ and K^+^ excretion (Figure 4). Similarly, pregnancy-induced adaptations in key distal tubule transporters (ENaC, ROMK, and H^+^-K^+^-ATPase) are required to prevent excess natriuresis and kaliuresis (Figure 5).
- Model simulations show that inhibition and knockout of the ENaC results in a notable increase in Na^+^ excretion and decrease in K^+^ excretion (Figure 6). This is consistent with experimental studies (see Ref. (41)).
- H^+^-K^+^-ATPase knockout mice exhibit altered pregnancy-induced adaptations in NCC, Pendrin, and ENaC that differ from pregnant wild type mice (see Table 2; Ref. (40)). Model simulation suggest that those altered adaptations serve to limit severe kaliuresis, at the expense of elevated Na^+^ excretion (Figure 7).
- Hypertensive female rats exhibit a similar shift as virgin females in Na^+^ transport from the proximal tubules to the distal tubules during pregnancy (Figure 8; Figure 9).

### Kidney function during pregnancy

The kidneys play a key role in maintaining homeostasis. Pregnancy is unique in that plasma volume expansion and electrolyte retention is required to sufficiently supply the rapidly developing fetus and placenta. Failure to expand plasma volume can lead to fetal growth restrictions (3,53–55). Gestational disorders such as gestational hypertension and preeclampsia have also been linked with insufficient plasma volume expansion (3,54–56). Indeed, renal adaptations must occur to be able to support such a significant change in the maternal body.

The proximal tubule is the first nephron segment and reabsorbs most of the filtered load, which is massively increased from the start of pregnancy (26,27,31). During pregnancy, proximal tubule length is increased leading to more transport capacity (27,34,57). Without this change, our models predict that excess natriuresis and kaliuresis will occur (Figure 4). While NHE3 activity has not been determined in pregnant rats experimentally, our simulation results indicate that without NHE3 upregulation, proximal tubule Na^+^ reabsorption is insufficient, resulting in increased urinary Na^+^ excretion (see Figure 4). Along the distal segments, the ENaC is greatly upregulated throughout pregnancy (41,48). Without this adaptation, model simulations reveal that excess natriuresis and K^+^ reabsorption will occur (Figure 5).

Several kaliuretic factors act on the kidneys of a pregnant rat: elevated aldosterone levels (3,58,59), decreased NCC activity (47), and increased ENaC activity (41,48,59). However, through pregnancy the maternal body is able to maintain K^+^ homeostasis and even retain K^+^ in late pregnancy (26,28). The K^+^ secretory channels in the distal segments, which in a virgin rat vigorously secrete K^+^, are downregulated (28). In addition, the active K^+^ transporter, H^+^-K^+^-ATPase has been shown to be significantly upregulated (28,29,40,60). Our model simulations show how together these transporter adaptations prevent excess K^+^ loss (Figure 5).

Given the large amount of fluid and solutes handled by the kidneys, minute adjustments in transport are sufficient to yield a difference in intake and urinary excretion. Synchronization of pregnancy-induced renal adaptations are required to sustain the retention of electrolytes and volume characteristic of pregnancy.

### Effect of distal tubule transporter inhibition on pregnancy

West et al. (41) found that chronic renal ENaC blockade (both pharmacological and genetic inhibition) ablated Na^+^ retention in pregnant rats leading to significant fetal growth restriction with a decrease in maternal blood pressure and body weight. Our model simulations on renal Na^+^ transport with both 70% ENaC inhibition as well as full ENaC knockout (i.e., 100% inhibition) predict a massive increase in Na^+^ urinary excretion in both mid- and late pregnancy due to decreased Na^+^ reabsorption along the distal convoluted tubule and connecting tubule (Figure 6A1; Figure 6A2). Additionally, secretion of K^+^ along the distal segments is reduced, ultimately leading to a decrease in urinary K^+^ excretion (Figure 6B1; Figure 6B2). This could lead to excess K^+^ reabsorption, especially in MP where K^+^ retention does not occur normally (26,28). Excess K^+^ reabsorption without other balancing mechanisms would lead to hyperkalemia (22,23,25). The model predictions presented in this study together with the experimental results from West et al. (41) emphasize the importance of the ENaC during pregnancy for both Na^+^ and K^+^ handling.

Walter et al. (40) studied the effects of pregnancy on H^+^-K^+^-ATPase type 2 knockout mice. They found that pregnant H^+^-K^+^-ATPase type 2 knockout mice not only had altered K^+^ balance but also had significantly inhibited plasma volume expansion (40). We note that while this is a rat model, adaptations that happen during pregnancy in rats and mice are similar, so we conducted simulations in a similar way here. Additionally, we note that no currently published study has investigated knockout of H^+^-K^+^-ATPase in rats during pregnancy.

We conducted two types of H^+^-K^+^-ATPase knockout simulations: one with full knockout of the H^+^-K^+^-ATPase transporter only (denoted by HKA-KO) and the second with H^+^-K^+^-ATPase knockout with the additional altered transporter adaptations in pregnant H^+^-K^+^-ATPase knockout mice reported in Walter et al. (40) (denoted by HKA-KO-preg; see *Materials and Methods* and Table 2). The HKA-KO simulations resulted in a higher increase in urinary K^+^ excretion than the HKA-KO-preg simulations when compared to the respective baseline models for both MP and LP (Figure 7B1; Figure 7B2). A similar change occurred in Na^+^ handling; HKA-KO simulations resulted in only a small increase in urinary Na^+^ excretion from baseline MP and LP model predictions, while the HKA-KO-preg simulations, which included a lower increase in ENaC activity, had an increase in urinary Na^+^ excretion (Figure 7A1; Figure 7A2).

During normal pregnancy, ENaC activity is massively upregulated from virgin values (26,41,48,59). Walter et al. (40) observed a lower increase in ENaC activity in pregnant H^+^-K^+^-ATPase type 2 knockout mice when compared to the pregnant wild-type mice. Our model results reveal that having lower ENaC activity during pregnancy in H^+^-K^+^-ATPase type 2 knockout mice there is less K^+^ secretion resulting in lower urinary K^+^ excretion (Figure 7B1; Figure 7B2). However, decreasing ENaC activity from normal pregnant levels also comes at the cost lower Na^+^ reabsorption (Figure 7A1; Figure 7A2). Since pregnancy-induced plasma volume expansion is driven by Na^+^ retention, it appears that the observed loss in plasma volume expansion (see Ref. (40)) in pregnant H^+^-K^+^-ATPase type 2 knock out mice may be driven by the compensatory changes in the nephrons. Specifically, the altered Na^+^ transporter adaptations from normal pregnancy values in the pregnancy-specific H^+^-K^+^-ATPase type 2 knockout mice (captured in HKA-KO-preg simulations; see Table 2) may work to prevent excess kaliuresis at the cost of losing more Na^+^ and hence resulting in lower plasma volume expansion that is known to be driven by Na^+^ retention.

Female rats, pregnant or not, express more Na^+^ transporters along the distal nephron segments compared to males (39,61,62). While the proximal tubule reabsorbs the bulk of the volume and electrolytes, the distal segments are responsible for “fine-tuning” tubular transport. To adapt to changes with the body, fine-tuning of electrolyte and volume reabsorption can best be achieved via signalling to the distal segments. Placing a larger transport load on the distal segments enables the females to adapt to the changing electrolyte and volume homeostasis requirements in pregnancy and lactation. When distal segment transporters are inhibited, or pregnancy-induced adaptations do not occur properly, our model simulations show that major alterations in electrolyte and volume renal handling will occur.

### Hypertension in females and pregnancy

Hypertension is a highly complex disease with many underlying pathophysiological mechanisms, some of which remain to be fully understood. Hypertension affects over 30% of the global adult population and is the leading cause of cardiovascular disease (12). It is estimated that the prevalence of hypertension will continue to rise due to ageing populations, increases in the exposure to high Na^+^ and low K^+^ diets, and a lack of physical activity (12).

During hypertension, inhibited pressure natriuresis reduces NHE3 activity and proximal tubular Na^+^ reabsorption, leading to a higher Na^+^ load to the distal segments (35,50,51). Interestingly, the medullary and cortical thick ascending limbs exhibit differential regulation of NKCC2, with the transporter down- and up-regulated in these two segments, respectively (35,37) (see Table 3). Simulation results obtained for the hypertensive virgin and mid-pregnant female models indicate that this differential regulation of NKCC2 minimizes Na^+^ retention while preserving K^+^ (Figure 9). This result is consistent with predictions from a previous modeling study of angiotensin II-induced hypertension in the male rat (see Ref. (36)). Differential regulation of NKCC2 in angiotensin II-induced hypertension may be attributed to angiotensin II stimulation of NKCC2 in the cortex but not in the medulla (35). Along the distal segments, the activity of the NCC and ENaC are both upregulated in hypertension (see Table 3). In both the hypertensive virgin and hypertensive MP models, this change led to increased Na^+^ reabsorption in the distal segments.

In hypertension, both with and without pregnancy, there is a shift in Na ^+^ transport from the proximal tubules to the distal tubules (see Figures 8 and 9). Given that the proximal tubules are more efficient than distal tubules in transporting Na^+^ (63), in that less oxygen is consumed to reabsorb a given amount of Na ^+^, the downstream shift implies an increase in overall oxygen demand. The elevated metabolic demand, if accompanied by maternal endothelial dysfunction that negatively impacts renal oxygen supply, may lead to renal hypoxia and kidney damage.

### Future work

Studying hypertensive disorders of pregnancy is complicated in that there are multiple types: chronic hypertension, chronic hypertension with superimposed pre-eclampsia, gestational hypertension, and pre-eclampsia (8,64). In this study we chose to investigate chronic hypertension due to little experimental data on nephron function in gestational hypertension and pre-eclampsia. Abreu et al. (49) investigated only mRNA data on a few transporters in hypertensive and normotensive pregnant rats. Hu et al. (65) investigated altered expression of renal Na^+^ transporters using urinary vesicles in pregnant women with pre-eclampsia. Other studies on renal transporter function during gestational hypertension or pre-eclampsia were not found. The changes we studied were primarily from studies that used angiotensin II-induced hypertension experimental protocol (35,37). Other experimental protocols have not reported as detailed transporter profiles during hypertension to date. As more experimental results are published, similar models to the ones presented in this study may be developed to understand renal function during these pregnancy-specific disorders.

Another common complication of pregnancy that likely affects the kidney is gestational diabetes (6). Gestational diabetes is diagnosed by hyperglycemia that develops in the third trimester (of human pregnancy) and resolves post-parturition. Jiang et al. (66) investigated the impact of high-fat diet induced gestational diabetes on the SGLT2 and GLUT2 transporters in mice nephrons. The present model can be modified to simulate gestational diabetes and its functional implications on kidney function as more experimental research is conducted.

In this study we focused on the impact of pregnancy and hypertension on renal function by using detailed cell-based epithelial transport models. Indeed, hypertension and pregnancy are both physiological states that are multi-factorial in nature. Specifically, increased blood pressure in hypertension is caused by a combination of impacts of renal handling, hormone systems, sympathetic nervous activity, and the cardiovascular system (67–69). No known whole-body pregnancy-specific blood pressure model exists to date (2). Indeed, existing models may be extended to consider this unique physiological state as well as the pathophysiology of hypertension. Similarly, since we are only modeling kidney function, we do not capture full body electrolyte homeostasis, considering the impacts of all organs involved in homeostasis. Models on electrolyte homeostasis may be extended to consider pregnancy to capture this electrolyte retention (2,23,70).

As average maternal age increases and the rates of comorbidities such as hypertension and diabetes increase in the population of women of childbearing age, rates of pregnancies complicated by a disorder will increase (6,7,10). However, our understanding of pregnancy physiology and especially gestational disorders remains limited. The study of pregnancy and its related pathologies has been historically limited by the fetal risks and ethical implications of running clinical trials on pregnant women (2,71). However, this limitation may be overcome, in part, by using computational models to test hypotheses, unravel complicated mechanisms, and potentially test the efficacy for therapies for gestational diseases *in silico* with the use of pregnancy-specific models as developed in this study. Indeed, there is much more computational work that can be done in understanding pregnancy physiology as well as gestational diseases as we recently reviewed in Ref. (2).

## Limitations of the study

Epithelial transport models have been extensively developed to provide an accurate accounting of solute and water transport to yield insights along with experimental evidence for the complicated transport pathways, driving forces, and coupling mechanisms that are involved in kidney function. These models have been used to study the functional implications of sexual dimorphisms (39,62,72–76), interspecies differences (75), diabetes (74,77), hypertension (36), hyperkalemia (78), circadian rhythms (76,79), and others. However, there are limitations that often stem from the paucity of experimental data as well as model structure as discussed in Refs. (42,80,81).

One key limitation is that in the models presented in this study it is assumed that the interstitial fluid composition is known a priori. There are changes in these models that may affect interstitial fluid composition, but since there is no evidence that has measured these changes, we did not include those changes in our model assumptions. Interactions among nephron segments and the renal vasculature can be modelled using the approaches in Refs. (82–85).

Similar to the limitations in our previous study (27), gestational kidney physiology has not been fully studied as there are some parameters that had to be assumed rather than directly based on experimental evidence. A similar limitation is there for the hypertension models. We used experimental evidence from Ref. (37) to inform our model parameter choices, however this study only investigated Na^+^ transporters. It is likely that there are other renal alterations that are impacted by hypertension. As more experimental evidence on renal transporter function during pregnancy and hypertension becomes available these models can be improved.

