## Supplementary Materials for "Effect of pregnancy and hypertension on kidney function in female rats: Modeling and functional implications"

### Supplemental Materials and Methods

***Juxtamedullary vs. superficial nephrons.*** The length of the long descending limbs and ascending limbs are determined by which type of juxtamedullary nephron is being modeled. To capture this, in the model there are six classes of nephrons: a superficial nephron (denoted by "SF") and five juxtamedullary nephrons that are assumed to reach depths of 1, 2, 3, 4, and 5 mm (denoted by "JM-1", "JM-2", "JM-3", "JM-4", and "JM-5", respectively) based on the length of the long descending limb. As derived in Ref. (1), the ratios for the six nephron classes are n_SF=2/3, n_(JM-1)=0.4/3, n_(JM-2)=0.3/3, n_(JM-3)=0.15/3, n_(JM-4)=0.1/3, and n_(JM-5)=0.05/3 so that 2/3 of the nephrons are superficial. Note that shorter juxtamedullary nephrons are the most common, thus turning in the upper portion of the inner medulla. All other segments in the juxtamedullary nephrons are the same as the superficial nephrons except the length of the cortical thick ascending limb and the connecting tubule. Since the glomeruli of juxtamedullary nephrons are located lower in the cortex than the superficial nephron glomeruli, these segments do not have to be as long for the nephron to pass the glomerulus at the macula densa. Hence, the cortical thick ascending limb and connecting tubule are modeled with a shorter length.

Additionally, it has been shown that the SNGFR for juxtamedullary nephrons is higher than the superficial nephron SNGFR (1,2). We assume that the juxtamedullary SNGFR is about 40% higher than the superficial SNGFR as in the virgin model (3).

The connecting tubules of the five juxtamedullary nephron types and the superficial nephron coalesce into the cortical collecting duct. To model this, we compute the flows from the six nephrons at the start of the collecting duct. The remaining model is the collecting duct which does not have distinct nephron segments. See Ref. (1) for more details on multi-nephron model development.

***Pregnancy-specific models.*** We created pregnancy-specific models to simulate kidney function in mid-pregnancy (MP) and late pregnancy (LP) by using the virgin (female-specific) multiple nephron epithelial transport model developed in Ref. (3) and increasing or decreasing relevant virgin model parameter values based on experimental findings in the literature. For changes in transporter activities for the MP and LP models we follow the same approach as in our previous study (Ref. (4)). These changes are briefly described below and discussed in more details in Ref. (4). Because transporter activity changes in pregnancy are driven primarily by hormonal adaptations, we assumed that the pregnancy-induced changes in transporter activity levels in the juxtamedullary nephrons are the same as the superficial nephrons.

In pregnancy, kidney volume increases (5,6). In particular, the proximal tubule, the first segment along the nephron where most $Na^+$ and $K^+$ reabsorption occurs, lengthens. Thus, we increased the proximal tubule length in the MP and LP models based on existing experimental measurements from Ref. (6,7). We also assume an increase in the diameter along the nephron, based on observed dilation in the collecting ducts during pregnancy (8). Without assuming a small tubular dilation, the much-elevated volume flow induced by the increased filtration would cause an excessive drop in tubular fluid pressure. Together increased proximal tubule length and nephron diameter result in a larger kidney volume.

We used experimental data to determine changes in parameter values for $Na^+$ transporters in our MP and LP models. Much of these results are reviewed in Ref. (9). Mahaney et al. (10) demonstrated region-specific changes in $Na^+$-$K^+$-ATPase activity and expression during pregnancy: $Na^+$-$K^+$-ATPase activity decreased throughout pregnancy in the cortex, but in the medulla, activity increase in MP while remaining unchanged in LP compared to virgin control (10). Recently, West et al. (11) reported that the activity of $Na^+$-$K^+$-$2Cl^-$ cotransporter 2 (NKCC2), a key transporter along the thick ascending limb, was significantly increased in LP and only slightly in MP. In the distal segments, the epithelial $Na^+$ channel (ENaC) fine tunes the remaining $Na^+$ in the luminal fluid before being excreted into urine. West et al. (12) reported that ENaC activity is nearly doubled in both MP and LP. Later findings showed renal adaptations in the ENaC are essential for sufficient $Na^+$ retention during pregnancy (13). In another study, West et al. (14) reported that the activity of the $Na^+$-$Cl^-$ cotransporter (NCC), which is found in the distal segments, is largely unchanged during MP and decreased during LP. Expression of the Cl-/bicarbonate exchanger, pendrin, in the connecting tubule and cortical collecting duct, is increased through pregnancy (15).

In the proximal tubule, the $Na^+$/$H^+$ exchanger (NHE3) drives much of the $Na^+$ reabsorption. However, $Na^+$/$H^+$ exchanger activity has not been well characterized during pregnancy. We note that it has been shown that in female rats (i.e., virgin), there is higher protein expression but lower activity of NHE3 when compared with male rats, indicating reserve NHE3 that can be activated during pregnancy (3,16,17). To avoid excess natriuresis, kaliuresis, and diuresis during pregnancy, we increased NHE3 activity in the MP and LP models based on the assumption that the reserve NHE3 in female rats is activated during pregnancy (see **Error! Reference source not found.**). An analysis of this assumption is discussed further in the Results and Discussion segments.

Aquaporin 2 (AQP2), the water channel in the collecting duct, is upregulated during MP and LP (18–20). It has also been shown that the water channel in the descending limb, AQP1, is upregulated during LP, but not significantly changed during MP (19). Based on these findings, water permeability in relevant segments was modified for increased AQP1 and AQP1 (see **Error! Reference source not found.**).

Since $K^+$ retention starts during LP, changes in $K^+$-specific renal transporters have mainly been studied during LP (9,21). The $K^+$-secretory channels in the distal segments, namely, the renal outer medullary $K^+$ channel (ROMK) and large-conductance $K^+$ channel (BK), are significantly downregulated, while $H^+$-$K^+$-ATPase pump activity is substantially increased during LP (21). We increased $H^+$-$K^+$-ATPase activity and decreased $K^+$ apical permeability in the appropriate distal segments in the LP model accordingly (see **Error! Reference source not found.**). Additionally, we hypothesized that the $K^+$-$Cl^-$ cotransporter in the ascending limb and distal convoluted tubule is upregulated during pregnancy to avoid excessive kaliuresis and natriuresis. While $H^+$-$K^+$-ATPase activity or BK permeability during MP remains poorly characterized, we note that Abreu et al. (18) showed that mRNA expression of ROMK2 is massively downregulated during MP. To avoid excessive kaliuresis and natriuresis in the MP model, we assume that similar changes to the $K^+$ transporters occur during MP.
